# A single cell spatial temporal atlas of skeletal muscle reveals cellular neighborhoods that orchestrate regeneration and become disrupted in aging

**DOI:** 10.1101/2022.06.10.494732

**Authors:** Yu Xin Wang, Colin A. Holbrook, James N. Hamilton, Jasmin Garoussian, Mohsen Afshar, Shiqi Su, Christian M. Schürch, Michael Y. Lee, Yury Goltsev, Anshul Kundaje, Garry P. Nolan, Helen M. Blau

## Abstract

Our mobility requires muscle regeneration throughout life. Yet our knowledge of the interplay of cell types required to rebuild injured muscle is lacking, because most single cell assays require tissue dissociation. Here we use multiplexed spatial proteomics and neural network analyses to resolve a single cell spatiotemporal atlas of 34 cell types during muscle regeneration and aging. This atlas maps interactions of immune, fibrogenic, vascular, nerve, and myogenic cells at sites of injury in relation to tissue architecture and extracellular matrix. Spatial pseudotime mapping reveals sequential cellular neighborhoods that mediate repair and a nodal role for immune cells. We confirm this role by macrophage depletion, which triggers formation of aberrant neighborhoods that obstruct repair. In aging, immune dysregulation is chronic, cellular neighborhoods are disrupted, and an autoimmune response is evident at sites of denervation. Our findings highlight the spatial cellular ecosystem that orchestrates muscle regeneration, and is altered in aging.

**Highlights:**

- Single cell resolution spatial atlas resolves a cellular ecosystem of 34 cell types in multicellular neighborhoods that mediate efficient skeletal muscle repair
- Highly multiplexed spatial proteomics, neural network and machine learning uncovers temporal dynamics in the spatial crosstalk between immune, fibrogenic, vascular, nerve, and muscle stem cells and myofibers during regeneration
- Spatial pseudotime mapping reveals coherent formation of multicellular neighborhoods during efficacious repair and the nodal role of immune cells in coordinating muscle repair
- In aged muscle, cellular neighborhoods are disrupted by a chronically inflamed state and autoimmunity

---

Our longevity depends on the renewal of tissues to meet the challenges of daily physical and molecular stresses. The repair process is critical for maintaining tissue function throughout life. Thus, a better understanding of regulatory mechanisms that operate during regeneration and the dysregulation that results from aging offer significant potential for the design of targeted therapies to enhance tissue repair and function (Blau and Daley, 2019; Blau et al., 2015; Fuchs and Blau, 2020).

Skeletal muscle accounts for ∼40% of our body mass and is subject to the physical stress of movement. Each muscle group consists of aligned contractile myofibers which are innervated by motor neurons and attach to bone via tendons. Skeletal muscle tissue is highly vascularized due to the metabolic demand of muscle contractions. In most scenarios, muscle damage incurred during exercise or injury such as muscle strains is efficiently repaired, and contractile function is restored. The repair of myofibers is carried out by muscle stem cells (MuSCs), also known as satellite cells (Blau et al., 2015; Relaix and Zammit, 2012; Wang and Rudnicki, 2012). MuSCs remain dormant in a quiescent state and respond to injury by proliferating to generate a pool of myogenic progenitors which fuse to form new myofibers (Dumont et al., 2015). However, in aging, muscles atrophy. This leads to molecular dysregulation that disrupts the signals that instruct MuSCs to proliferate and orchestrate the complex cellular symphony that underlies the regenerative process, resulting in fatty-fibrotic scarring and progressive replacement of muscle cells (Blau et al., 2015; Mann et al., 2011; Muñoz-Cánoves et al., 2020). This regenerative deficit exacerbates the muscle loss seen with aging.

The niche, or stem cell microenvironment, is a critical determinant of the regenerative response (Fuchs and Blau, 2020). Single cell analysis (De Micheli et al., 2020a; Giordani et al., 2019; Porpiglia et al., 2017) and genetic ablation approaches (reviewed in Bentzinger et al., 2013a; Fuchs and Blau, 2020) have suggested the requirement for coordinated interactions between cell types to carry out repair. The yin and yang function of immune cells is highlighted by their critical role in normal repair, and their disruption of muscle function in inflammatory myopathies, dystrophies, and aging. When transient, proinflammatory signals and macrophage recruitment initiate the wound-healing response and activate MuSCs. This process is carefully regulated, as persistent immune responses in muscles afflicted with muscular dystrophy and systemic changes in inflammatory cells and cytokines in advanced age, through a process termed “inflammaging" (Ferrucci and Fabbri, 2018), are associated with progressive fibrotic accumulation and progressive loss of muscle function. These studies suggest that cells reside in a delicate regenerative ecosystem in which complementary, interconnected, and interdependent relationships with other cell types are essential to carry out their programmed function in rebuilding the tissue.

Despite this knowledge, there are major gaps in our understanding of the ecosystem underlying the process of regeneration and of aging, largely due to limitations in currently used technologies. For example, cell-cell interactions cannot be assessed by methods that require tissue dissociation, such as flow cytometry, CyTOF, or single cell RNA-sequencing. Critical information is lost, for instance about changes to the niche, a microenvironment in which spatially localized cell-cell signaling and extracellular matrix (ECM) interactions are key to efficacious regeneration. Spatially restricted regulators likely determine cell migration behavior and fate (Bentzinger et al., 2013; Blau et al., 2015; Fuchs and Blau, 2020; Wang and Rudnicki, 2012). On the other hand, traditional histological methods, in which cell integrity within tissues is maintained intact, suffer from the limited capability of visualizing only 3-4 proteins simultaneously due to secondary antibody cross-reactivity and spectral overlap. Thus, most currently used methods fail to reveal the complexity of spatially localized interactions of diverse cell types, the ECM, and secreted molecules that mediate regenerative regulatory mechanisms.

Here we overcome this limitation by employing multiplex imaging to simultaneously profile the spatial distribution of cell surface, intracellular and ECM proteins during skeletal muscle regeneration and aging in mice. We explore the interplay of the plethora of cell types that spring into action to restore the complex architecture of skeletal muscle tissues after injury. We developed analytic tools that utilize neural networks to identify tissue features and unsupervised clustering for identifying 34 cell types at single cell resolution to build a single cell spatial atlas of muscle regeneration and of aging, tools that will serve as a resource for similar studies in other tissues. We uncover positional information and the temporal dynamics of intercellular crosstalk between immune, fibrogenic, vascular, nerve, and myogenic cells at sites of injury and repair, and their relationship to the extracellular matrix in multicellular neighborhoods. We employ methods we developed for spatial pseudotime mapping to build a regeneration clock of cell interactions and how they change over time, an unbiased metric of the repair process. This analysis uncovers a nodal role for immune cells in efficacious muscle regeneration and in the disruption of the cellular ecosystem that accompanies aging. Finally, our atlas not only provides single cell resolution tissue architecture of skeletal muscle and a holistic overview of cell-cell interactions that underly muscle repair and aging, but also provides a roadmap for using neural network and unsupervised clustering approaches to understand complex changes in cellular neighborhoods that underly biological processes in a wide range of tissues.

## Results

### Identification and validation of a skeletal muscle cell regeneration antibody panel

We aimed to create a comprehensive atlas detailing how distinct cell subtypes contribute to muscle regeneration after injury. To date, our knowledge of the temporal dynamics of the various cell types that participate in muscle repair derives from techniques that dissociate the tissue like flow cytometry, CyTOF, and single cell RNAseq (Bentzinger et al., 2013; De Micheli et al., 2020b, 2020a; Giordani et al., 2019; Petrany et al., 2020a; Porpiglia et al., 2017), which lack information regarding spatial relationships and cell-cell interactions. Histological studies that retain the tissue intact have suffered from limitations due to the inability to simultaneously visualize more than ∼4 markers simultaneously due to spectral overlap of fluorophores using immunofluorescence. While current spatial transcriptomics approaches offer insights into cellular relationships in situ, they are currently limited by low resolutions (∼10-50um) that cannot truly resolve single cells and lack the ability to resolve how cells interact with the ECM. Here we overcame these limitations by using CO-Detection by indEXing (CODEX; Fig. 1B), a high resolution method that allows up to 60 protein markers to be visualized simultaneously by iterative probe binding and microscopy in a single tissue section (Goltsev et al., 2018; Kennedy-Darling et al., 2021; Schürch et al., 2020). As a result, the diverse cell types involved in efficacious regeneration can be definitively identified. This is achieved by a combination of multiplex imaging and localized protein profiling which resolves the temporal progression of the various cell subtypes and their spatial organization that are inherent to efficacious skeletal muscle regeneration.

**Figure 1.**
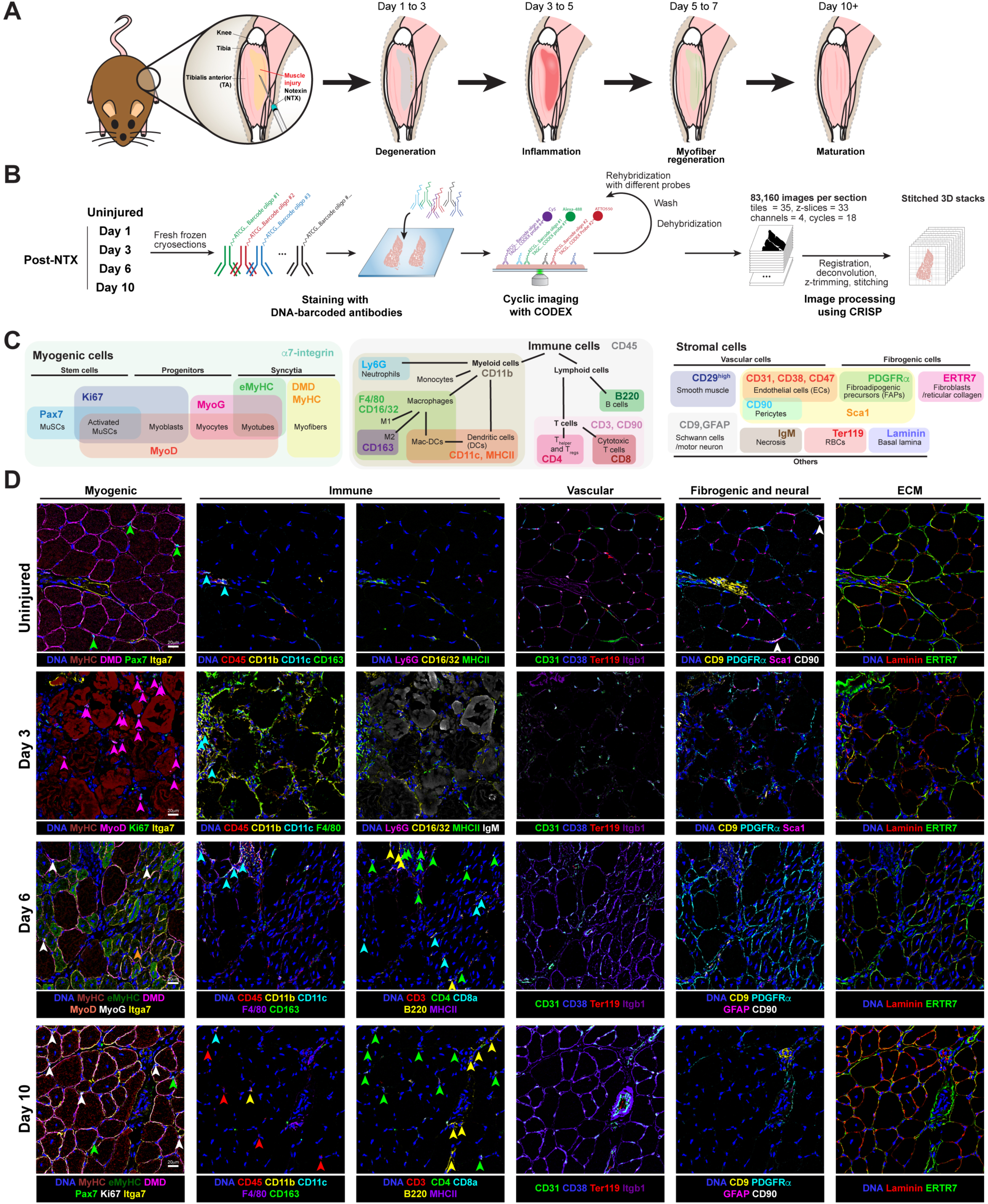
Multiplexed immunofluorescence imaging to elucidate cellular heterogeneity during skeletal muscle regeneration. **A)** Schematic of myotoxin-induced murine skeletal muscle injury and regeneration timeline. The tibialis anterior (TA) muscles of young and aged mice were injected intramuscularly with notexin to induce myofiber damage and regeneration. **B)** Schematic of multiplexed imaging of regenerating skeletal muscle tissues using CODEX. Muscle tissues were cryosectioned onto coverslips, stained with a panel of DNA barcoded antibodies, and rendered by cyclic imaging with fluorophore conjugated cDNA probes using CODEX. Multicycle tissue images were registered, deconvolved, trimmed, and stitched using CRISP image processor. **C)** Antibody panel design to resolve cell types found during skeletal muscle regeneration. Overlapping and mutually exclusive protein markers were used to distinguish biologically relevant cell types and subsets. **D)** Representative CODEX images of uninjured and regenerating muscle sections. Pseudo-colored antibody staining as indicated below each image. The same field-of-view is shown across each time point (row). Markers of cell types within each lineage are shown in each column.

### Spatial profiling of regenerating skeletal muscle by multiplex imaging

We induced muscle injuries by intramuscular injections of notexin (NTX), a well characterized myotoxin that causes local myofiber damage, and monitored changes in cell-cell relationships using CODEX in transverse tissue sections over a 10-day time course during which the injury is repaired (Fig. 1A). NTX damage models a grade 2 muscle strain that occurs in sports or traumatic injuries, where muscle tears but does not undergo complete rupture (Pollock et al., 2014). To construct a CODEX antibody panel that encompasses all cell lineages, we combined previously reported cell type-specific markers (Bentzinger et al., 2013) with additional markers identified by single cell analysis of muscle (De Micheli et al., 2020a; Giordani et al., 2019; Porpiglia et al., 2017). This panel identifies myogenic, immune, vascular, fibrogenic, and motor neuron cells and their functional subsets (Fig. 1C). To distinguish a progression of myogenic cell states, we used established markers: MuSCs (Pax7), proliferating myoblasts (MyoD and Ki67), committed myocytes (myogenin (MyoG)), myotubes (embryonic myosin heavy chain; eMyHC) and myofibers (dystrophin (DMD) and adult myosin heavy chain (MyHC)) (Bentzinger et al., 2012; Silberstein et al., 1986). We used well characterized markers to distinguish immune cell types of the myeloid lineage: monocytes (CD11b), neutrophils and granulocytes (CD11b and Ly6G), macrophages (CD11b and F4/80) and dendritic cells (CD11b, CD11c, and class II major histocompatibility complex (MHC-II)) (De Micheli et al., 2020a; Giordani et al., 2019). We further distinguished macrophages by expression of CD16/32 (FcR) on FcR+ macrophages (Fitzer-Attas et al., 2000) and CD163 on M2 macrophages (Hu et al., 2017) that distinguish these subsets from M1 macrophages. To differentiate immune cell types of the lymphoid lineage we used established markers for B cells (B220) and T cells (CD3 and CD90) (Bendall et al., 2011). Within the T cell population, CD4 was used to identify T Helper and T Regulatory cells, and distinguish them from CD8 marked cytotoxic T cells. We were also able to identify multiple vascular cell subtypes in the muscle tissue, including endothelial cells (CD31 and Sca1) and smooth muscle cells (a7-integrin (a7-int) and b1-integrin (CD29)). We used CD9 to mark the Schwann cells surrounding the motor neurons that innervate muscle tissue (Anton et al., 1995). To capture the fibrosis that is a feature of damaged tissue, we identified fibroadipogenic progenitors (FAPs) by their expression of PDGFRα and Sca1 (Joe et al., 2010; Uezumi et al., 2010). Tenocytes comprise the tendons that connect the muscles to the bones and were identified by tenomodulin (TNMD) (Docheva et al., 2005; Giordani et al., 2019). Finally, we visualized the tissue ECM that provides structural and biochemical support to the tissue (laminin and reticular collagen (ERTR7)). To mark the regions of damage, we included IgM which has been shown to bind to damaged, necrotic myofibers (Petrany et al., 2020b). We validated our antibody panel by ensuring that cell subtypes co-expressed multiple markers (e.g., co-staining of macrophages by CD45, CD11b and F4/80) and that distinct cell type subsets were clearly identifiable based on detection of unique markers (e.g., pericytes were distinguished by CD90 expression). Multiplexed detection of this array of antibodies allowed us to resolve specific from non-specific signal that can be detected by certain antibodies (as described in the limitations section) and discern temporal changes in antibody intensity and localization (Fig. S1). Together, this spectrum of antibodies enabled resolution of the dynamic alterations in the abundance and organization of various cell types throughout the regeneration time course.

### Deep learning to map regenerating muscle

Multiplex imaging benefits from sub-micrometer resolution (20x magnification; 0.377um/pixel), but generates massive amounts of data per experiment. To analyze this large dataset, we developed a set of computational tools to register and stitch images obtained from automated microscopes across imaging cycles, identify and segment single cells from the stitched images, and classify identified cells based on antibody staining (Fig. 2A and S2).

**Figure 2.**
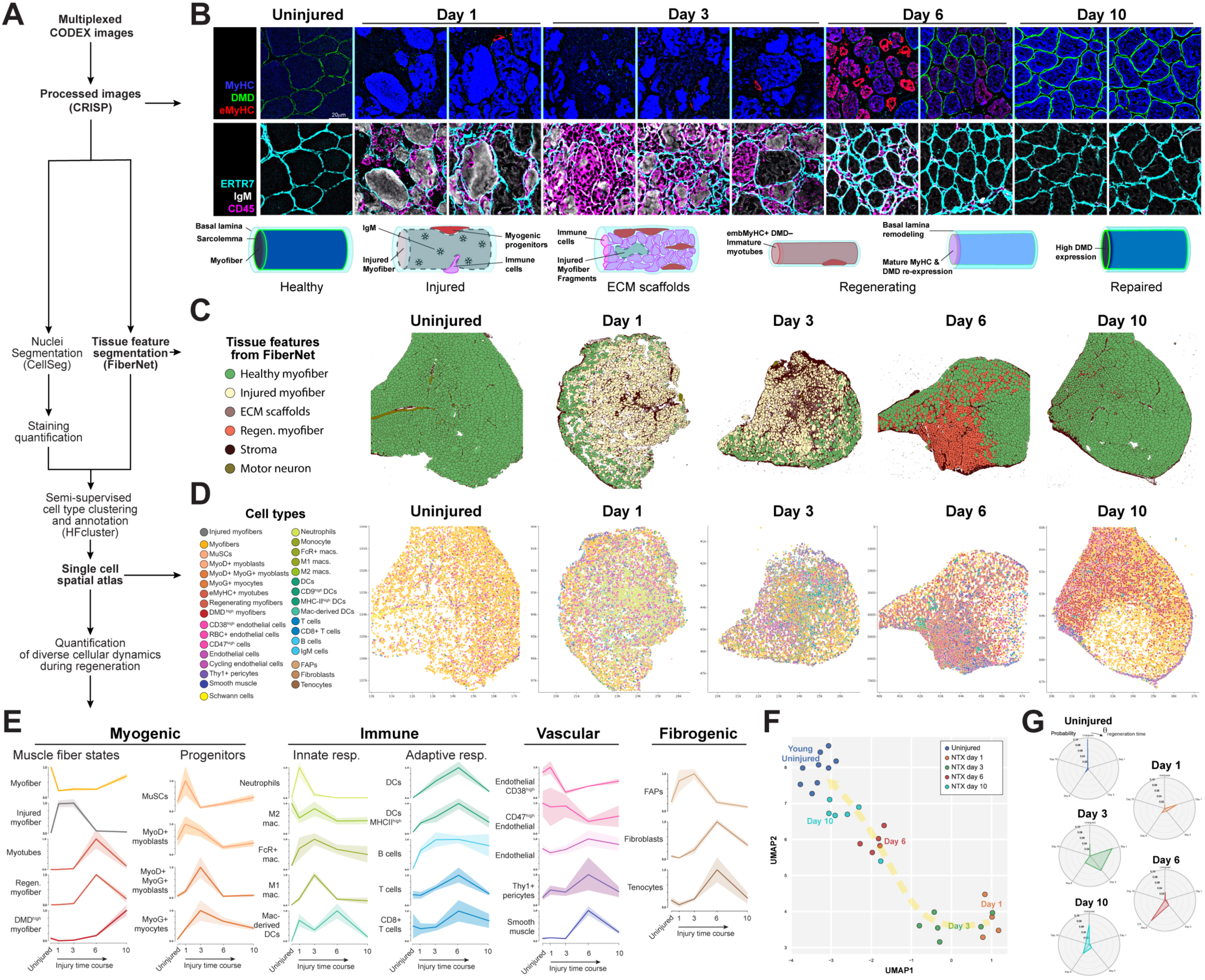
Single cell spatial atlas of skeletal muscle regeneration. **A)** Schematic of computational analysis pipeline to resolve spatial relationships from multiplexed imaging data. **B)** CODEX images of myofiber states in uninjured muscles and in a time course after injury. Healthy muscle fibers express myosin heavy chains (MyHC, Blue) and dystrophin (DMD, green) on their sarcolemma, and are surrounded by the endomysium marked by ERTR7 (cyan). The myotoxin used induces sarcolemmal damage, resulting in the loss of DMD and leads to the accumulation of IgM (grays) in the injured myofibers. Immune cells marked by CD45 (magenta) infiltrate the muscle at days 1 and 3. Myogenic progenitors differentiate and express embryonic isoforms of myosin Myh3 (eMyHC; red) marking newly formed myotubes. By day 6, eMyHC expression was reduced in regenerating myofibers that mature and begin to re-express DMD. By day 10, the muscle structure are largely restored but the regenerated myofibers showed higher DMD expression. The same field-of-view is shown in each column. A cartoon representation of each stage of myofiber degeneration, regeneration is shown below each respective panel. **C)** Representative FiberNet classification of skeletal muscle fiber states and stromal regions based on multiplexed imaging data. Images are pseudo-colored by the classification labels of tissue features from FiberNet according to the legend (left) **D)** Representative cell type annotation of uninjured and regenerating skeletal muscles regions based on multiplexed imaging data. Each dot is one nucleus; Prospectively annotated cell type is represented by the color in the legend (left). **E)** Temporal enrichment of cell types during skeletal muscle regeneration. Enrichment is min-max normalized for each cell type. Error bands represent s.e.m. n=4-8 per time point. **F)** UMAP embedding of the cellular composition of uninjured and regenerating skeletal muscles. Arrow indicates the regeneration trajectory from day 1 after injury to the uninjured state. **G)** Temporal variance of cell types found in muscles at each regeneration time point. Polar coordinates represent the regeneration time course and probability distribution of cells found in each time point across all regeneration time points.

We developed the CRISP image processing pipeline (Palla et al., 2021) to align and register our CODEX images in 3D at sub-pixel resolution. The improved image alignment and stitching CRISP provides enhanced our ability to perform in silico tissue clearing (remove autofluorescence signal) and reduced imaging artifacts. Once our images were registered, stitched, and cleared, we leveraged the exceptional image recognition abilities of convolutional neural networks (CNNs) to segment cells and tissue features. We used CellSeg, a CNN trained for the segmentation of nuclei (Lee et al., 2022), to generate masks of each nucleus and quantify the intensity of staining of each antibody within the nuclear and perinuclear compartments (Fig. S3A).

While CellSeg allowed us to characterize many of the features of our tissue, its reliance on single nuclei data complicates its use in analyzing the large multinucleated myofibers and ECM structures characteristic of muscle tissue. To solve this issue, we created FiberNet, a CNN trained to recognize muscle fiber states (healthy, injured, and regenerating myofibers) and features marked by ECM (ECM scaffolds, motor neurons, and stroma) in fluorescence images (Fig. 2B). We defined “healthy muscle fibers” as those that expressed mature MyHC and dystrophin (DMD). By contrast, “injured” myofibers (labelled by IgM) exhibited a loss of DMD and a7-int due to the destructive effect of notexin on the sarcolemma. The ECM remained intact, providing a reference of the location of the myofiber prior to damage. This provided a scaffold-like structure encompass a range of cell types including eMyHC+ differentiating myotubes resided after IgM+ injured myofibers were removed. As regenerating myofibers matured, they decreased expression of eMyHC and began expressing DMD. The ECM was significantly thicker around regenerating myofibers and exhibited increased staining for ERTR7. In addition, ECM markers identify complex structures such as connective tissue of stromal regions and motor neuron tracts. Remarkably, FiberNet identifies each of these features with ∼98% accuracy (Fig 2C).

### Cellular heterogeneity and dynamics of skeletal muscle regeneration

We first used previously charted notexin injury regeneration timecourses to validate our time course of regeneration and progression of myofiber states (Bentzinger et al., 2013; Hardy et al., 2016; Morton et al., 2019). Quantification of muscle cross sections after injury revealed that 1 day after injury ∼80-90% of muscle fibers were damaged (Fig. S3B). ECM scaffolds devoid of IgM+ injured myofiber debris were transiently detected at day 3 before regenerating myofibers formed at day 6. By day 10 post injury, newly formed myofibers increased in size and returned to a healthy state.

Using CODEX we were able to gain an in-depth view of the dynamic changes of cell subsets and their interplay during regeneration after injury at a single cell resolution (Fig. 2B). We sought to classify single cells in the tissue into cell types based on their antibody staining patterns. The automated identification of cell types from imaging data remains a computational challenge. Unlike high-throughput sequencing approaches, multiplex imaging approaches have lower dimensionality, suffer from imaging artifacts, and exhibit variable tissue autofluorescence all of which contribute to poor performance in clustering algorithms. To overcome these limitations, we developed a high-fidelity clustering pipeline (HFcluster) that is optimized for multiplexed imaging and immunofluorescence data. HFcluster is unique in that it overcomes the contribution of non-specific signals and other noise while clustering by first learning potential cell types in the tissue using the antibody staining patterns of cells with robust signal and then propagating those identities onto cells with lower signal that share similar staining patterns (Fig. S3C). We further improved our clustering by integrating tissue feature classifications from our FiberNet algorithm, which facilitated the classification of myofiber states. Using this approach, we identified 34 distinct cell subsets that matched the expected combination of markers defined by our selected panel of antibodies (Fig. S4; additional details in Supplementary Methods). Clustering results were consistent across tissues, time points and experimental batches (Fig. S4). Since each cell was indexed with its spatial coordinates, we were able to generate a single cell resolution atlas of skeletal muscle regeneration which includes the positional information for each of the 34 distinct cellular subsets throughout the time course of muscle regeneration (Fig. 2D). A progression from intact myogenic cells was followed on day 2 of injury by a dramatic influx of immune cells, followed on day 3 by an increase in endothelial and vascular cell types, which increased on day 6 and resolved on day 10 as muscle regeneration nears completion. Interestingly, while FiberNet classified regenerated myofibers at day 10 as healthy, many of these fibers showed higher DMD expression than in uninjured muscle, thus suggesting longer lasting molecular differences in regenerated myofibers (Fig. 2C-D).

We identified several endothelial cell (EC) subsets distinguished by marker expression and spatial localization. We detected a range of expression of CD38, Sca1, and CD47 on other ECs (Fig. S4). Interestingly, our data reveal that CD38, a cell surface nicotinamide adenine dinucleotide nucleosidase, specifically marks capillary ECs but not the ECs of larger blood vessels (Fig. 1D, S4 and S5A). After injury, expression of CD38 in capillary ECs correlated with the presence of the red blood cell marker Ter119, suggesting a relationship with capillary perfusion (Fig. S4, S5B-C). CD38+ ECs markedly decline in the injured areas of day 3 muscles (Fig. 2E). By day 6 and 10, CD38+ capillary ECs are found in regions with DMD^high^ myofibers but not in regions that continue to express embryonic myosins (Fig S5B-C). These findings suggest that CD38 expression in ECs is restricted to perfused capillaries, revealing previously uncharted changes to tissue perfusion and angiogenesis during late stages of muscle repair.

We quantified each cell type subset across the regeneration time course to discern the temporal dynamics of changes in cell composition that occur during regeneration (Fig. 2E). Our analysis revealed that as muscle tissues transition through regeneration, there is a continuous flux of functional subsets of myogenic, immune, vascular and fibrogenic cellular lineages. Cellular composition is distinct at each time point (Fig. S4), and matches previously established dynamics of myogenic differentiation, and innate and adaptive immune responses as quantified by methods entailing tissue dissociation (Bentzinger et al., 2013; De Micheli et al., 2020a; Giordani et al., 2019; Porpiglia et al., 2017; Tidball, 2017).

Since the abundance of specific cell subsets is in constant flux (Fig. 2E), we sought to capture the transient states of tissue regeneration through the composition of cells in each tissue and determine the overlap of subsets at each time point. Using our single-cell cell type data from CODEX and Uniform Manifold Approximation and Projection (UMAP), a dimension reduction technique (McInnes et al., 2020), we assessed the compositional similarity of each tissue. This analysis revealed that regeneration time points can be distinguished by the relative abundance of cell subsets within each tissue (Fig. 2F). We found that cell types in uninjured, day 1, day 3, and day 6 samples were largely non-overlapping (Fig. 2G). Day 3 tissues contained the largest diversity of subsets and shared common cell types with day 1 and day 6 tissues (Fig. 2G). Day 10 tissues contained the most subsets in common with uninjured tissues but still contained regenerating cell types in common with day 6 tissues (Fig. 2G). These data outline a temporal cellular composition regeneration trajectory that culminates in a near return to an uninjured state (Fig. 2F). They also provide insights into the cell type composition at the tissue level that can be used to gauge the regeneration status of the muscle. Together, these findings establish a high resolution temporal spatial atlas of muscle regeneration and suggest distinct temporally determined cellular functions.

### Defining cellular neighborhoods of regenerating muscle

We noted that some cellular subtypes from distinct cellular lineages displayed correlated dynamics, which led us to postulate that they co-exist in cellular neighborhoods. For example, there is an inverse correlation of cell types between healthy myofibers and injured myofibers (Fig. 3B). Thus, spatial relationships between pairs of cell types are often directional, and an enrichment of cell types in the vicinity of each other suggests grouping or dispersion dynamics, and that inter-lineage regulation between these distinct cell subtypes can occur. This finding fits well with findings by others that cells are known to organize into cellular neighborhoods through chemoattractant or chemorepellent signals (Goltsev et al., 2018; Schürch et al., 2020) and reside in niches that depend on the presence or absence of other cells (Fuchs and Blau, 2020).

**Figure 3.**
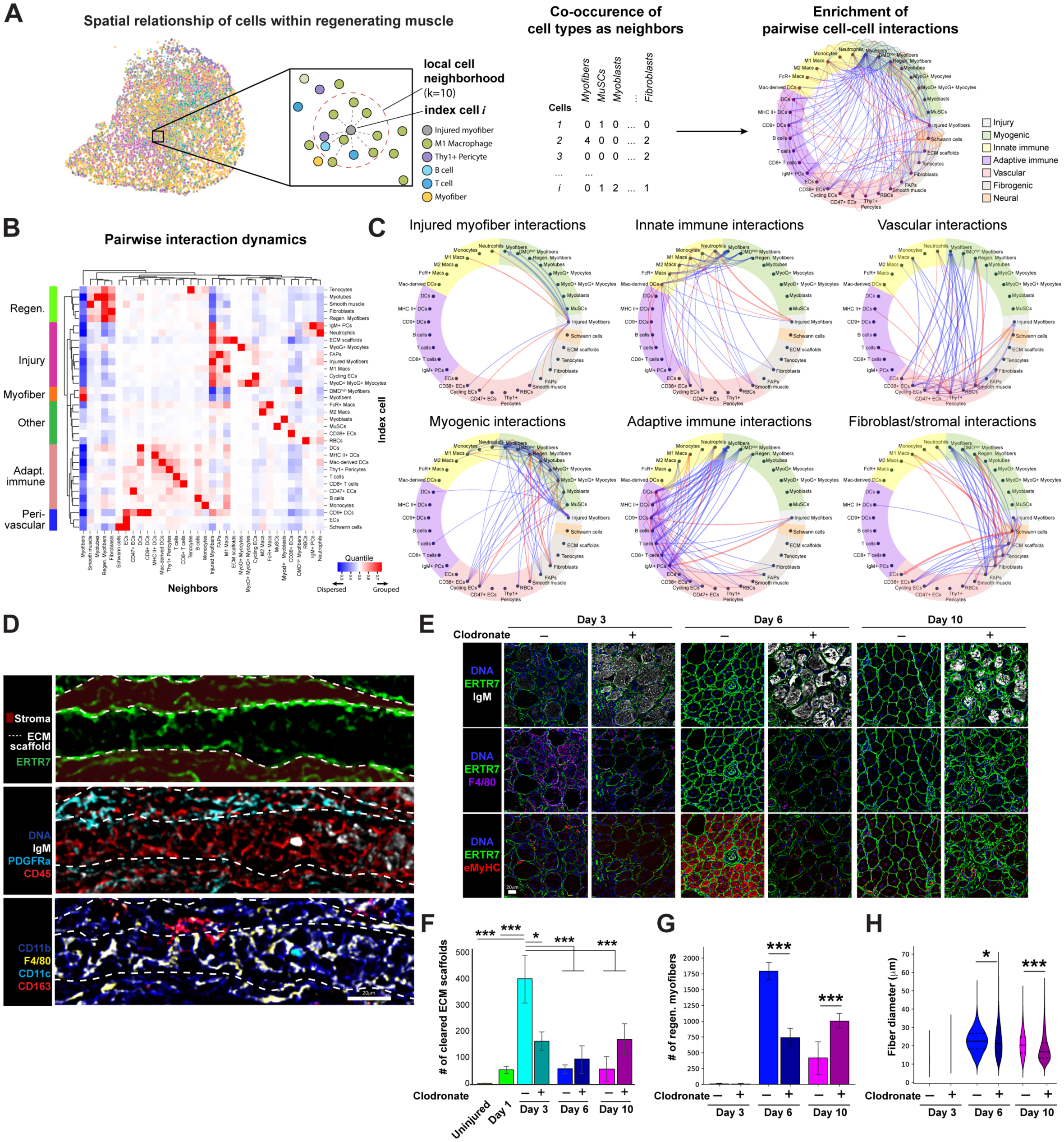
Spatial interactions among regenerative cell types of skeletal muscle. **A)** Schematic of single cell spatial analysis to identify enrichment pairwise interactions between cell types. Index cells and their nearest neighbors were quantified, and the co-occurrence of cell types in proximity was used to identify enriched interactions. **B)** Heatmap of pairwise cell-cell interactions during skeletal muscle regeneration. Positive enrichment (red) represents cell type pairs that were found in proximity at rates more than expected; Negative enrichment (blue) represents cell type pairs that were found in proximity at rates less than expected. Hierarchical clustering identified correlations between cell type pairs that represents co-interactions that could be grouped according to biological processes occurring during regeneration (left). **C)** Cross lineage interactions during skeletal muscle regeneration. Enrichment of pairwise interactions is indicated by arrows. Cell types are arranged by cell lineages as indicated in Fig. 3A. Arrows indicate direction of spatial dynamics; arrow thickness is indicative of enrichment. Red arrows indicate grouped (>0.55 quantile) and blue arrows indicate dispersed dynamics (<0.45 quantile) . **D)** Longitudinal views of extracellular matrix (ECM) scaffolds and infiltrating cell types around injured myofibers at day 3 after myotoxin injury. ECM scaffolds (dashed lines) were marked by ERTR7 (green, top panel); IgM+ injured myofibers (grays), PGDFRa+ FAPs (cyan), CD45+ immune cells (red) and DAPI (blue) shown in the middle panel; CD11b+ myeloid cells (blue), F4/80+ macrophages (yellow), CD11c+ dendritic cells (cyan), and CD163+ M2 macrophages shown in the bottom panel. The same field-of-view is shown across all panels. **E)** Representative images of regenerating muscles at day 3, 6, and 10 after injury with or without intramuscular injection with clodronate liposomes at day 2. IgM+ injured myofibers (grays, top panels); F4/80+ macrophages (magenta, middle panels); eMyHC+ myotubes and regenerating myofibers (red, bottom panels); ECM scaffolds were marked by ERTR7 (green); DAPI (blue). The same field-of-view is shown in each column. **F)** Quantification of ECM scaffolds in regenerating muscles at day 3, 6, and 10 after injury with or without intramuscular injection with clodronate liposomes at day 2. **G)** Quantification of regenerating myofibers in regenerating muscles at day 3, 6, and 10 after injury with or without intramuscular injection with clodronate liposomes at day 2. **H)** Minimum axis lengths of regenerating myofibers in regenerating muscles at day 3, 6, and 10 after injury with or without intramuscular injection with clodronate liposomes at day 2. **(F-H)** Error bars represent s.e.m.; n=4-8 per group; * p<0.05; ** p<0.01; *** p<0.005.

To gain insights into the spatial arrangement of cell types in cellular neighborhoods of regenerating muscle, we analyzed the co-occurrence of cell subsets (neighbors), quantifying grouping and dispersion relationships between cell type pairs (Fig. 3A). We defined the largest set of cell-cell interactions around injured myofibers (Fig. 3B). Pairwise interaction analysis revealed clusters of spatially enriched cell types that correspond to regenerative processes (Fig. 3B), including clusters of interactions driven by injured myofibers, adaptive immune cells, vasculature, regenerating myotubes, and healthy myofibers, as well as a cluster of other interactions involving M2 macrophages, MuSCs and CD38+ ECs.

Injured myofibers exhibited reciprocal attractive relationships with early inflammatory cell types including neutrophils and M1 macrophages (Fig. 3C, top left). Cycling ECs, FAPs, and myogenic progenitors (myoblasts and myocytes) were also enriched in the vicinity of injured muscle fibers, suggesting that factors released at the site of injury may facilitate the cell cycle re-entry of muscle resident stem cells (Fig. 3C, top left). Consistent with a previous report (Verma et al., 2018), we identified an enrichment of CD38+ capillary ECs in the vicinity of MuSCs, suggesting that these cells are a part of the MuSC niche. However, the enrichment is unidirectional, MuSCs are not enriched in the vicinity of CD38+ capillary ECs, indicating CD38+ ECs do not require MuSCs in their niche (Fig. 3B-C).

In other neighborhoods, we uncovered changes in the interaction of immune, vascular, and stromal in response to injured myofibers, as well as temporally distinct supportive cell types that co-occur with subsets of myogenic cells in neighborhoods in which new myofibers are being formed. Neutrophils and macrophages (M1 and FcR+ subsets) mount an innate immune response to injury (Fig. 3C, top middle). These myeloid subsets largely associate with each other, recruit monocytes, and interact with dendritic cells (Fig. 3C, top middle). While most infiltrating immune cells are myeloid in accordance with an innate immune response, the accumulation of IgM in injured myofibers is consistent with an antibody mediated adaptive immune response. Indeed, injured myofibers were enriched in neighborhoods comprised of a subset of CD9+ dendritic cells and IgM+ plasma cells (Fig. 3C, bottom middle). Dendritic cells and CD9+ dendritic cells interacted with B cells and T cells in lymphoid aggregates that form around regenerating myofibers at day 6 after injury (Fig. 3C, bottom middle). Myogenic progenitors (myoblasts and myocytes) are associated with M1 macrophages and cycling endothelial cells typical of an early regenerative state, whereas fused myotubes and regenerating myofibers are associated with fibroblasts, tenocytes, and smooth muscle cells (Fig. 3C, bottom left and right) characteristic of a later regenerative state.

A common feature of a later regenerative phase and uninjured muscle is that among immune cell interactions is that they all show repulsion dynamics with regenerating myofibers and mature myofibers (DMD^high^ and healthy subsets) (Fig. 3C, middle panels), indicating that the presence of mature myofibers suppresses inflammatory cell types. Similarly, most vascular and fibrogenic cell subsets exhibit repulsion dynamics with myofibers except for CD38+ ECs (Fig. 3C, top and bottom right). This finding underscores the known association of capillary CD38+ ECs intertwined with the myofibers in the muscle vasculature. It also highlights the known anti-fibrotic effects of myofibers on FAP differentiation (Joe et al., 2010; Uezumi et al., 2010; Wosczyna et al., 2019). Additionally, our analysis established that motor neuron-associated Schwann cell neighborhoods are enriched in M2 macrophages and accompanying vessels consisting of ECs (CD38–), which supply the nerve with nutrients (Fig. 3C, bottom right). These cell-cell interaction dynamics point to coordinated temporally regulated cellular interactions that occur in series during muscle regeneration. Specifically, injury triggers inflammation and stem cell activation, that in turn recruits additional cell types, which promote differentiation in a coordinated cascade of events entailing precisely orchestrated changes in cellular neighborhoods.

### Cells that traverse the myofiber basal lamina during regeneration

ECM structures like the basal lamina can act as barriers, allowing only select cell types to traverse them. Since ECM scaffolds are comprised of structural proteins that are destroyed upon dissociation, the ability of cell types to traverse the ECM scaffold has remained elusive. While such scaffolds have been visualized previously by electron microscopic and intravital imaging (Vracko and Benditt, 1972; Webster et al., 2016), CODEX imaging allows us to capture the heterogeneous population of cells within these ECM scaffolds during regeneration (Fig. 2A, 3D, and S6A). In longitudinal sections along the length of the muscle, CODEX reveals ECM scaffolds as tracts outlined by precisely aligned reticular collagen fibrils (stained by ERTR7; Fig 3D, top; and S6A) which accumulate IgM+ debris from injured myofibers replete with infiltrating CD45+ immune cells (Fig. 3D, middle). This contrasts with the localization of PDGFRa+ FAPs, which are mostly found outside the ECM scaffolds (Fig. 3D, middle), presumably because the ECM acts as a barrier to these cells.

To gain further insights into cellular heterogeneity within ECM scaffolds and relationships between cell types that traverse the ECM, we quantified cell subsets found in ECM scaffolds and assessed their co-occurrence at each stage of regeneration. Cells were mapped based on their spatial location relative to the ECM scaffold, enumerated and identified by FiberNet, then clustered by similarity of cellular composition (Fig. S6B). As expected, M1 macrophages were the major cell type found within ECM scaffolds. M1 macrophages were associated with small numbers of other cell types such as monocytes, other macrophage subsets, dendritic cells (DCs), fibroblasts, and regenerating myofibers (Fig. S6B-C). The differential localization of the myeloid cell population was further resolved based on marker expression (Fig. 3D, bottom). While CD11b+ myeloid cells were found both inside and outside of ECM scaffolds, many M1 macrophages (F4/80+ CD163–) but few CD163+ M2 macrophages were found inside the ECM scaffolds. F4/80+ CD11c+ cells were observed inside ECM scaffolds at day 6, suggesting a process of differentiation from macrophages to dendritic cells. Additional cell types such as Ly6G+ neutrophils, CD31 ECs, and myogenic progenitors were also found within ECM scaffolds at different time points (Fig. S6A). These results suggest that the ECM scaffold is at times a highly dynamic environment where cells readily migrate across the residual endomysium and basal lamina of the myofiber after injury.

Clustering analysis identified distinct ECM scaffolds that were either predominantly populated by myoblasts or by MuSCs and MyoG+ myocytes (Fig. S6C; clusters 27 vs. 21), consistent with contact mediated feedback on MuSC self-renewal from differentiating myocytes. Temporally, neutrophil-dominant and M1 macrophage-dominant ECM scaffolds appeared on day 1, became macrophage-dominant by day 3 and macrophage-derived DC-dominant by day 6 (Fig. S6D-E). Most other clusters that contained primarily non-macrophage cell types appeared at later regeneration time points after day 3, suggesting that M1 macrophages facilitate the transit of other cell types into the ECM (Fig. S6E).

### M1 macrophages clear the way for muscle repair

Macrophages play a critical role in tissue repair and signal to other support cells to coordinate their functions (Arnold et al., 2007; Brigitte et al., 2010; Chazaud et al., 2003; Du et al., 2017; Ratnayake et al., 2021; Shang et al., 2020; Tidball, 2017). Our pairwise interaction analysis indicated that M1 macrophages are enriched near ECM scaffolds (Fig. 3C, bottom right) and M1 macrophages are the predominant cell type that traverses ECM scaffolds (Fig. S6B). Moreover, the presence of M1 macrophages is largely mutually exclusive with other cells that traverse the ECM scaffolds, suggesting that M1 macrophage activity could be a rate limiting step. Thus, we hypothesized that a major function of macrophages within the ECM scaffold is to pave the way for myogenic cells to carry out regeneration.

To test this hypothesis, we performed intramuscular injections of clodronate liposomes to deplete macrophages locally in muscles at day 2 after injury and assessed regeneration dynamics by CODEX multiplex imaging (Fig 3E). While we were unable to deplete all M1 macrophages, the number of M1 macrophages within the myofiber basal lamina was significantly reduced. Consistent with this reduction, the number of M1 macrophage dominated ECM scaffolds was diminished on day 3 and cell type dynamics were aberrant (Fig. S6E-F). In accordance with our hypothesis that M1 macrophage traversal across the myofiber basal lamina is required to remove injured myofibers, clodronate treated samples contained IgM+ injured myofiber debris even at day 10 (Fig. 3E and S6A). M1 macrophages eventually infiltrated the muscle, however, regeneration was significantly delayed (Fig 3E and 3G). Clodronate treated muscles at day 6 had 50% fewer and smaller caliber regenerating myofibers (Fig. 3G). We also observed that myogenic differentiation was stalled, as there was an increase in ECM scaffolds containing MuSCs and myocytes at day 6 and fewer scaffolds contained mature regenerating myofibers at days 6 and 10 (Fig. S6F; clusters 21, 5 and 18, respectively). The regenerating myofibers found in clodronate treated muscles at day 10 were abnormal in their organization due to a persistence of myofiber debris post-injury which acted as a physical barrier preventing proper fusion of myocytes and constraining hypertrophic growth (Fig. 3E). These regenerating myofibers exhibited a 25% reduction in minimum diameter and did not reach a mature state, resulting in 60% fewer regenerated healthy muscle fibers by day 10 (Fig. 3G and S6B). Additionally, the loss of M1 macrophages allowed for an increase in traversal of fibroblasts across the basal lamina at day 6 and DCs at day 10 (Fig. S6E-F), suggesting that early M1 macrophage loss has a broad impact on the entire cellular response of the regenerating microenvironment.

### The M1 macrophage is a nodal regulator of regeneration

Having established that removal of M1 macrophages from the regenerating microenvironment caused significant delays in the repair process (Fig. 3E-H), we sought to characterize the extent and mechanism of the delay by quantifying the cell types present in the tissue across time to impute a “regeneration pseudotime”. Most of the cell subsets present during a normal regeneration time course appeared transiently. To this end, we encoded each cell in the tissue with the average timepoint at which it appeared. We calculated a local tissue pseudotime by averaging encoded times for each cell in specific tissue regions marked by a 75 x 75 μm grid (Fig. 4A,B). We compared this local regeneration pseudotimes with the post-injury time point of tissue collection (Fig. 4A,C). Using this approach, we were able to accurately distinguish injured and uninjured areas and predict the relative regeneration time of the injured regions (Fig. 4B). This pseudotime analysis also allowed us to visualize and quantify localized delays in regeneration at day 6 and 10 induced by macrophage depletion instigated by clodronate treatment of muscles (Fig. 4C).

**Figure 4.**
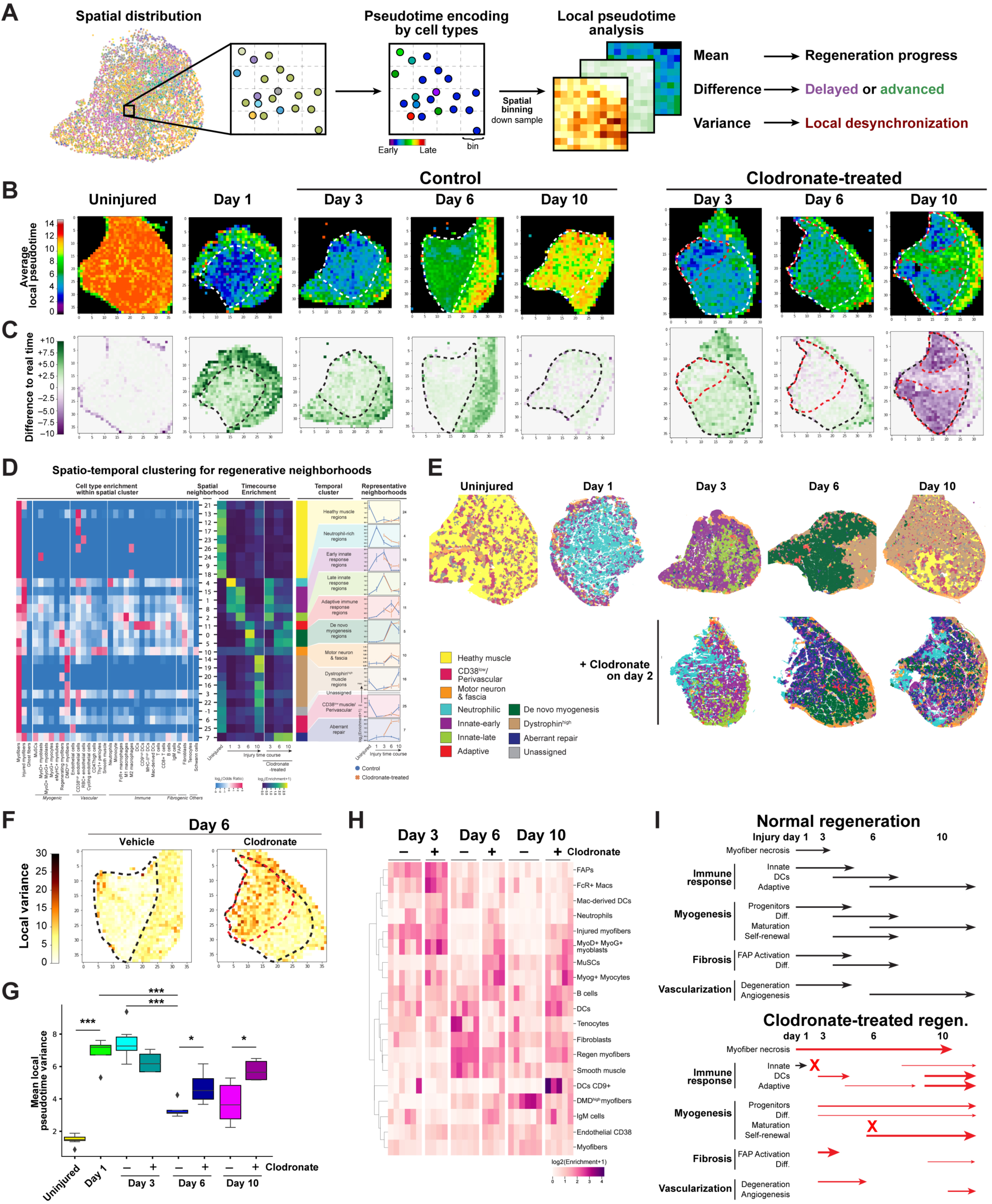
Spatial pseudotime and cell neighborhood analysis of tissue regeneration upon macrophage depletion. **A)** Schematics of spatial pseudotime analysis to reveal regeneration dynamics. Positional information of each cell is encoded with a pseudotime. Cells within a tissue can be sampled in a grid to estimate the mean local pseudotime, which can be compared with actual time after regeneration to estimate accelerated or delayed repair. High variance of cell pseudotimes in each grid space indicates the co-occurrence of cells that normally appear at different stages of regeneration, suggesting dysregulation or desynchronization of cellular processes. **B)** Mean local pseudotime of uninjured and regenerating muscles at day 1, 3, 6, and 10 after injury with or without intramuscular injection with clodronate liposomes at day 2. White dashed lines outline the injured region; red dashed lines outline regions affected by clodronate. **C)** Difference of local pseudotime to actual time points of uninjured and regenerating muscles at day 1, 3, 6, and 10 after injury with or without intramuscular injection with clodronate liposomes at day 2. Green and purple represent an accelerated or delayed regeneration, respectively; Black dashed lines outline the injured region; red dashed lines outline regions affected by clodronate. **D)** Spatial temporal cell neighborhood analysis of uninjured and regenerating muscles. Local cell compositions were clustered into spatial neighborhoods, revealing patterns of cellular interactions (left); spatial neighborhoods were further clustered by temporal dynamics during regeneration and after clodronate-treatment (middle heatmap) into temporal clusters. Temporal dynamics of representative spatial neighborhoods are shown for each cluster (right). Error bars represent s.e.m. of relative enrichment in control or clodronate-treated samples; n=4-8 per group. **E)** Representative temporal neighborhood clusters in uninjured and regenerating muscles at day 1, 3, 6, and 10 after injury with or without intramuscular injection with clodronate liposomes at day 2. Images are pseudo-colored by cell neighborhood clusters from panel **D**. **F)** Variance of local pseudotime of uninjured and regenerating muscles at day 6 after injury with or without intramuscular injection with clodronate liposomes at day 2. Increased local variance indicates the co-occurrence of cells that normally appear at different stages of regeneration; Black dashed lines outline the injured region; red dashed lines outline regions affected by clodronate. **G)** Quantification of local pseudotime variance in uninjured and regenerating muscles at day 1, 3, 6, and 10 after injury with or without intramuscular injection with clodronate liposomes at day 2. Boxes indicate mean, upper and lower quartile range; whiskers are 1.5 times the inter quartile range; n=4-8 per group; * p<0.05; ** p<0.01; *** p<0.005. **H)** Heatmap of cellular dysregulation in after macrophage depletion by intramuscular clodronate injection. Log transformed enrichment of cell types in regenerating muscles at day 3, 6, and 10 after injury with or without intramuscular injection with clodronate liposomes at day 2. Each column is a biological replicate; n=4-8 per group; Cell types showing significant change (p<0.05) with clodronate treatment are shown. **I)** Schematic of cellular dysregulation in after macrophage depletion by intramuscular clodronate injection. Arrow width indicates the relative alteration compared to normal regenerative conditions. X indicates a complete halt or absence of a given cell type.

We investigated whether the delay in regeneration after macrophage depletion could be due to a blockade in cellular progression through a normal regeneration program or via an alternative non-productive program. Since macrophage depletion by clodronate treatment disrupts myofiber regeneration and impacts a range of cell types that normally traverse the ECM (Fig 3G,H and S6F), we hypothesized that localized disruptions in key cell effectors like macrophages could further alter the spatial arrangements of cells and the timing of the normal regenerative program. To assess this, we performed a spatiotemporal cell neighborhood analysis, clustering the local co-occurrence of cell subsets during the regeneration time course. We identified 26 spatial neighborhood clusters with distinct cell type compositions (Fig. 4D). When we then cluster these spatial neighborhoods by temporal enrichment to determine which ones appear in a temporally regulated manner, we find 10 temporal clusters that change in abundance across the regeneration time course (Fig. 4D,E). These temporal clusters represent unique cell-cell interaction neighborhoods that occur during regeneration: healthy muscle, perivascular, neutrophilic infiltration, early and late innate immune response, adaptive immune response, and de novo myogenesis cell-cell interaction neighborhoods (Fig. 4D,E).

To determine if the delay in regeneration we observe after macrophage depletion arises from a block to the normal regeneration program or from an alternative non-productive program, we performed spatiotemporal clustering on our clodronate treated samples and compared cell-cell neighborhoods and their temporal appearance within the neighborhoods. If macrophage depletion merely delayed the normal regenerative process, we would expect the local compositions of cells after clodronate treatment to match those seen during normal regeneration. However, our cell neighborhood analysis revealed a cell neighborhood (spatial neighborhood 7) that was unique to clodronate treated samples (Fig. 4D,E). The aberrant neighborhood contained a disorganized conglomeration of injured myofibers, myogenic cell subsets characteristic of all stages, innate and adaptive immune cells, and fibroblasts (Fig. 4D,E), which normally appear in a precisely orchestrated sequence during normal regeneration (Fig. 4D). To quantify the extent of the deviation, we measured the pseudotime variance of cells in clodronate treated muscles within gridded tissue regions and compared our results to the pseudotime variance we determined during normal regeneration (Fig. 4A,F,G). We found that injured areas of normally regenerating muscle had low pseudotime variance, suggesting that normal regeneration is a temporally cohesive cellular process (Fig. 4F,G). In contrast, in clodronate treated samples pseudotime variance by day 6 and 10 was significantly increased. This is consistent with our cellular neighborhood analysis. These results suggest that macrophage depletion not only disrupts the progression of various cell types through the regeneration program, but also triggers the formation of aberrantly regenerating regions that contain cells that do not normally co-exist at the same time (Fig. 4F,G).

To gain an in-depth understanding of the aberrant regenerative process triggered by macrophage depletion, we assessed changes in the progression of specific immune, myogenic, vascular, and fibrogenic cell subsets in clodronate treated samples. We observed universal tissue disruption, marked by significant changes in cell subtype abundance across multiple lineages at all time points (Fig. 4H). In clodronate-treated muscles at day 3 after NTX, we observed increases in the numbers of neutrophils (p=0.03) that are normally resolved by day 3, yet also increases in FcR+ macrophages (p=0.01) and macrophage-derived DCs (p<0.01) that normally become abundant at day 6, providing evidence of a temporally aberrant accumulation of myeloid subsets (Fig. 4H). In addition, compared to untreated day 3 samples, the abundance of MyoD+ MyoG+ myoblasts and CD38+ ECs also increased by 2.4 (p<0.02) and 1.5-fold (p<0.02), respectively, indicating that macrophage depletion profoundly impacts the abundance of cells of other lineages (Fig. 4H).

These early changes coupled with the accumulation of dead myofiber debris that prevents proper myocyte fusion precipitates changes in additional cell types (Fig. 4H). We observed significant increases in DCs (p<0.005) and the appearance of a novel CD9+ DC population (p=0.001) by day 10. MuSC and MyoG+ myocyte abundance also increased at day 6 and 10 (p=0.001 and 0.0040; and p=0.0065 and 0.0226, respectively). Coupled with the sustained presence of regenerating myofibers (p=0.001) and a loss of DMD^high^ myofibers (p=0.001) at day 10, these findings are consistent with a delay in myotube maturation. Fibroblast abundance at day 6 (p=0.0043) was also decreased; this is consistent with our pairwise interaction analysis where fibrogenesis is coupled with myofiber maturation during regeneration.

Together, our data suggest that M1 macrophages play a pivotal role in coordinating regeneration. Upon their depletion by clodronate treatment, a block in phagocyte function creates a physical barrier to regeneration, which in turn triggers widespread disruption to the standard regeneration progression of immune, myogenic, fibrogenic, and vascular cell subtypes. These changes, in turn, disrupts regenerative cell neighborhoods, temporally desynchronizing regeneration, and accelerating adaptive immunity. As a result, MuSC function is shifted toward self-renewal, myofiber formation is hindered, and fibrosis and angiogenesis are delayed (Fig 4I).

### Aging changes skeletal muscle architecture

Aging is associated with numerous maladaptive changes in skeletal muscle including chronic inflammation, partial denervation, persistent fibrosis, and diminished regenerative capacity, which together lead to muscle wasting (Blau et al., 2015; Larsson et al., 2019; Muñoz-Cánoves et al., 2020). Such aging-associated effects have been probed at the transcriptomic level, both in bulk and at single cell resolution (Petrany et al., 2020a; Schaum et al., 2020; Tabula Muris Consortium, 2020), but how aging impacts the spatial organization of cells within muscle tissue has not been thoroughly explored. To address this, we performed CODEX multiplex imaging on skeletal muscle isolated from aged mice (25-28 months) and compared it to young muscle (2 mo) to understand the how the spatial cellular neighborhood composition of aged muscle changes and how this could lead to diminished regeneration.

Our regeneration pseudotime analysis of aged uninjured mouse muscles showed that the local cellular composition of aged muscles at steady state most closely resembles day 10 of young muscle regeneration (Fig. 5A,B). This regressed state is widespread throughout aged muscles and is in stark contrast with the localized degeneration and regeneration triggered by injury in young (Fig. 5A). We also noted an increase in the variance of regeneration pseudotimes of cells found in aged muscles, suggesting that aged muscles are more heterogeneous, concurrently containing cell subtypes normally found at distinct regeneration timepoints in young (Fig. 5C,D). This aberrant composition is apparent even when aged tissues are compared to day 10 of regeneration in young. We validated this observation by creating a UMAP projection of tissues representing our regeneration time course. Aged tissues clustered separately from day 10 regenerating muscles, suggesting that aging alters cell composition compared to uninjured or day 10 regenerating young muscle (Fig. S7A).

**Figure 5.**
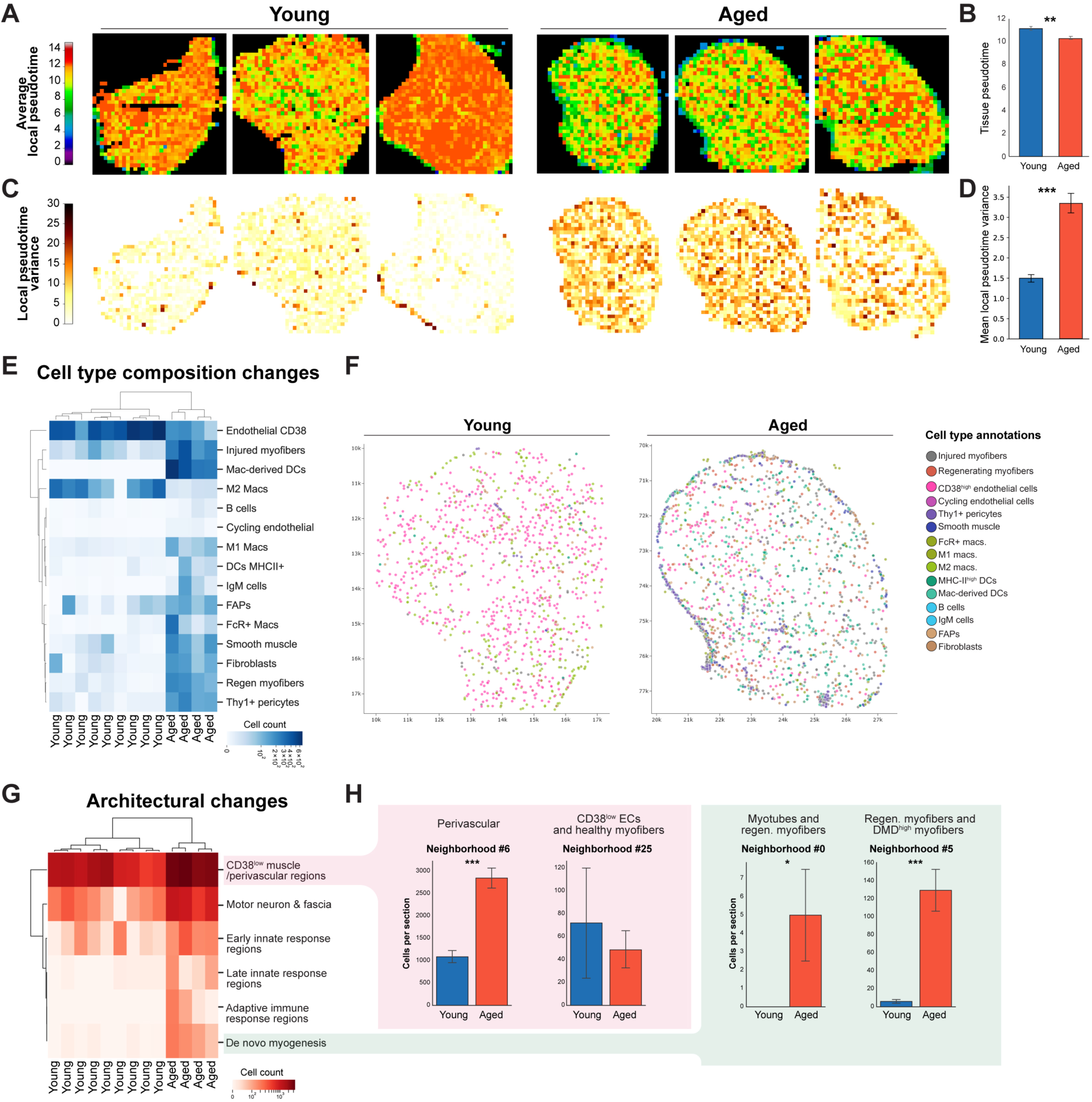
Localized cellular and architectural changes in skeletal muscle associated with murine aging. **A)** Mean local pseudotime of uninjured muscles of young and aged mice. **B)** Quantification of mean local pseudotime of uninjured muscles of young and aged mice. n=8 young and 4 aged samples; ** p<0.01. **C)** Variance of local pseudotime of uninjured muscles of young and aged mice. **D)** Quantification of local pseudotime variance of uninjured muscles of young and aged mice. n=8 young and 4 aged samples; *** p<0.005. **E)** Heatmap of cellular dysregulation in aged muscle. Log transformed enrichment of cell types in uninjured muscles of young and aged mice. Each column is a biological replicate; n=8 young and 4 aged samples; Cell types showing significant change (p<0.05) with aging are shown. **F)** Representative tissues showing spatial localization of dysregulated cell types in uninjured muscles of young and aged mice. Each dot is one nucleus; Prospectively annotated cell type is represented by the color in the legend (right). **G)** Heatmap of tissue architectural dysregulation in aged muscle. Log transformed enrichment of temporal neighborhood clusters from Figure 4D in uninjured muscles of young and aged mice. Each column is a biological replicate; n=8 young and 4 aged samples; Clusters showing significant change (p<0.05) with aging are shown. **H)** Expanded analysis of change in spatial neighborhood subclusters in uninjured muscles of young and aged mice. n=8 young and 4 aged samples; * p<0.05; *** p<0.005.

Uninjured aged muscles are enriched for cell types consistent with myofiber damage and repair, not young uninjured muscle (Fig. 5E), although the cell types are less abundant than we observed following young muscle injury (Fig. S4). CD38+ ECs and M2 macrophages found in the stroma of uninjured young muscles were less prevalent in aged muscles (p=0.004 and 0.02, respectively). We also observed an increase in the numbers of both injured myofibers (p<0.005) and regenerating myofibers (p=6x10^-6^). In addition, we detected cell composition changes associated with early injury and late regeneration simultaneously in aged tissues. FcR+ and M1 macrophages and FAPs that normally increase in numbers in early regeneration were more abundant in steady state uninjured aged muscles (p=0.015, 2x10^-6^ and 0.02, respectively). In conjunction, MHC-II+ DCs, macrophage-derived DCs, B cells, smooth muscle cells, Thy1+ pericytes, and fibroblasts that normally increase in late regeneration were also more abundant in aged muscles than in young (p=0.005, 2x10^-5^, 0.008, 3x10^-6^ and 0.0005, respectively). Thus, aged muscles are characterized by a persistently dysregulated regenerative state.

Importantly, several of the observed cell types (Thy1+ pericytes, macrophage-derived DCs) were aberrantly localized in the muscle fascia, the connective tissue encapsulating the muscle, and to small perivascular clusters within the muscle proper (Fig. 5F). The patterns observed were highly specific; a careful analysis of the perivascular/CD38^low^ clusters revealed that only the perivascular cluster, not the ECs and healthy myofiber cluster, was increased in aged muscle (Fig. 5H). Overall, consistent with our pseudotime and cell composition analysis, our neighborhood analysis revealed an increased abundance of clusters of innate and adaptive immune cell types, de novo myogenesis, motor neuron/fascia, and perivascular/CD38^low^ in aged muscle at steady state that is characteristic of the normal regeneration process seen in young (Fig. 5G, S7B).

Together, our data indicate that a combination of increased myofiber turnover, innate and adaptive inflammation, and vascular and fascia remodeling characterize aged muscles, which contribute to the loss of tissue homeostasis and alterations in the tissue microenvironment observed in aging. Moreover, our analysis reveals for the first time the complex microenvironmental changes at the molecular, cellular, and tissue architecture level that occur with aging.

### Autoimmunity in aged muscle

To probe molecular mechanisms that underlie the changes to the spatial neighborhood composition of aged muscle, we identified antibodies from our CODEX antibody panel that were differentially abundant in aged muscles compared to young (Fig 6A). We observed significant changes in myogenic markers (MyoD and MyoG) in myogenic neighborhoods, as well as immune markers (CD163, FcR, CD8a, CD11b) in inflammatory neighborhoods (Fig 6A). However, IgM was the sole factor that was differentially abundant across 6 of the 8 significantly changed neighborhoods we identified in aged muscle (Fig 6A).

**Figure 6.**
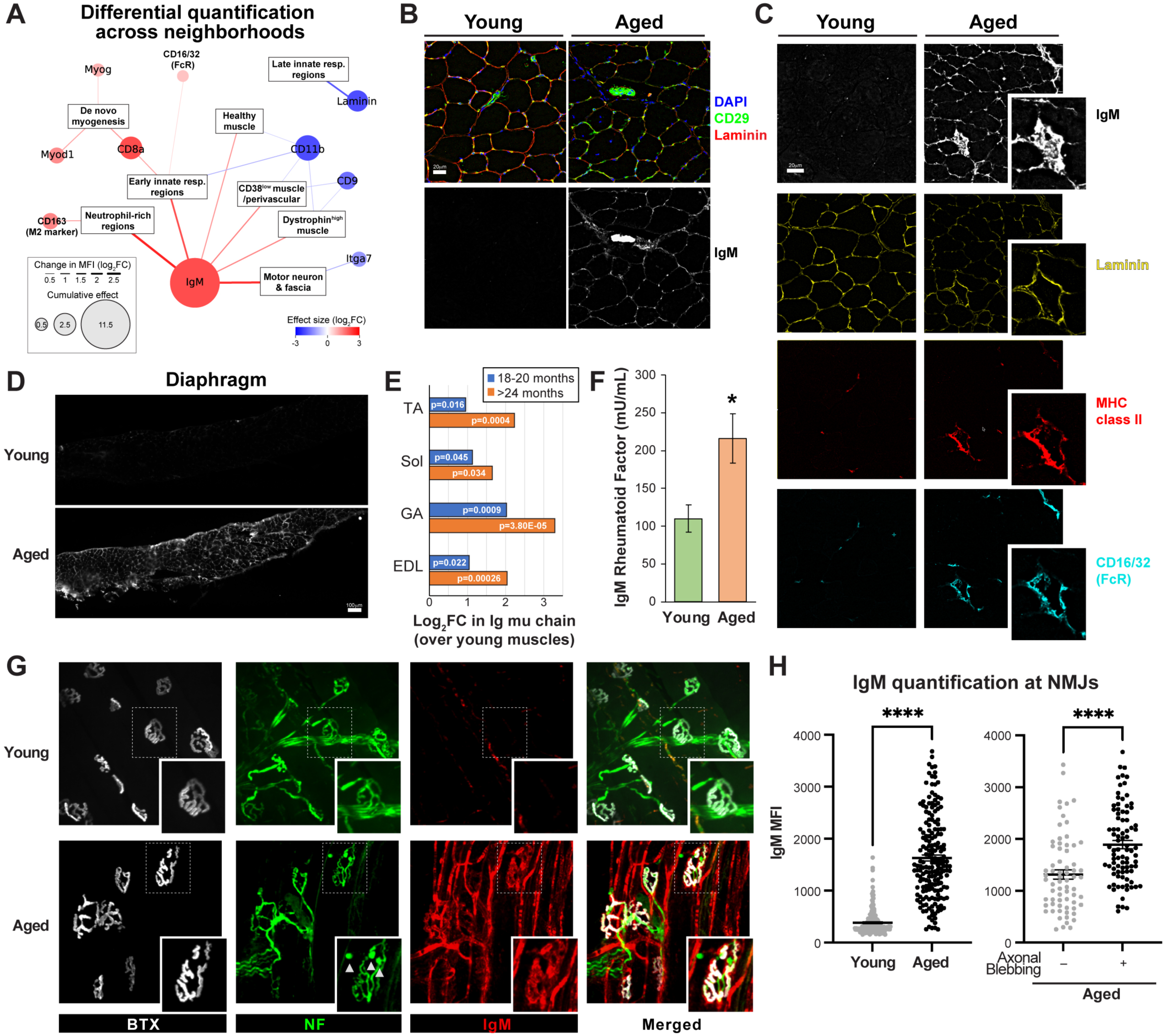
Age-related extracellular accumulation of IgM in murine skeletal muscle. **A)** Network representation of differential mean intensity analysis of CODEX images between uninjured muscles of young and aged mice. Lines represent significant change in staining intensity in the connected temporal neighborhood cluster. The size of circles for each marker indicated the cumulative effect across all connected neighborhoods. n=8 young and 4 aged samples; Markers showing significant change (p<0.05) with aging are shown. **B)** Representative CODEX images of uninjured muscles of young (left) and aged (right) mice. CD29 (green, top panels) marks myofiber sarcolemma and vasculature; Laminin marks the basal lamina (red, top panels); IgM staining (grays, bottom panels); DAPI (blue). The same field-of-view is shown in each column. **C)** Representative CODEX images of uninjured muscles of young (left) and aged (right) mice. IgM staining (grays, top panels); Laminin marks the basal lamina (yellow, 2^nd^ row panels); Major-histocompatibility class II molecules (MHC-II I-A/I-E, red) and Fc-gamma receptors (CD16/32, cyan) marks immune cells (3^rd^ and 4^th^ row panels); DAPI (blue). The same field-of-view is shown in each column. Insets show enlarged examples of IgM staining colocalized with immune markers. **D)** Representative traditional immunofluorescence histology for IgM in uninjured diaphragm muscles young (top) and aged (bottom) mice. **E)** Protein mass spectrometry quantification of the IgM mu chain in young and aged skeletal muscles, reanalysis of Schüler et al. 2021. Log fold-change over detected levels in muscles of young mice. **F)** IgM rheumatoid factor ELISA of serum from young and aged mice. n=4 young and 3 aged samples; * p<0.05. **G)** Representative immunofluorescence of neuromuscular junctions (NMJs) in wholemount uninjured EDL muscles from young (top) and aged (bottom) mice. Bungarotoxin (BTX, grays) marks acetylcholine receptors on the myofibers; neurofilament (NF; green) marks the motor neuron; IgM (red); merged image (right panels). Arrows indicate axonal blebbing observed in aged samples. Insets show enlarged examples of NMJs. **H)** Quantification of IgM staining intensity at neuromuscular junctions (NMJs) in wholemount uninjured EDL muscles from young and aged mice (left); and aged NMJs stratified by the appearance of axonal blebbing (right). n=3 young and 3 aged samples; Each dot is one NMJ. **** p<0.001.

IgM is the highest molecular weight immunoglobulin and is one of the first to appear upon antigen stimulation or microorganism exposure. Our data show that during regeneration IgM accumulates transiently in young injured myofibers, in conjunction with the influx of immune cells. Thus, we hypothesized that a persistent accumulation of IgM is a indication of autoimmunity in aged mice and could facilitate the aberrant proinflammatory changes we note in aged muscle.

Unique to aged muscle, we also observed strong IgM staining in the ECM and vasculature (Fig. 6B-C, S7C). IgM enriched areas in aged muscle also frequently harbored an abundance of immune cells that expressed FcR and MHC-II (Fig. 6C, S7C), which suggests that IgM promotes inflammation. We validated this finding across a range of muscle groups including diaphragm, tibialis anterior, gastrocnemius, and extensor digitorum longus muscles by traditional immunofluorescence using fluorophore-conjugated antibodies to mouse IgM (diaphragm data shown in Fig. 6D). To further understand the aging-associated changes in tissue IgM levels, we analyzed recently published proteomic mass spectrometry data of young and aged muscles (Schüler et al., 2021). We found that the mu chain of IgM is consistently detected as one of the top age-related proteomic changes, increasing over 8-fold in gastrocnemius muscles of aged mice (Fig. 6E). Overall, these data identify IgM as a systemic factor that accumulates in aged muscle.

It has previously been shown that IgM antibodies made in response to specific antigens can cross react with self-proteins including host IgG and act as rheumatoid factors (Dresser, 1978). Given the IgM accumulation we observed in the ECM of aged skeletal muscles, we hypothesize that this IgM could target self-proteins, contributing to the decline in muscle function characteristic of aging. To test this, we compared the autoreactivity of IgM isolated from the sera of young and aged mice to characterize its potential as a rheumatoid factor. Sera from aged mice showed a ∼2-fold increase in IgM rheumatoid factor compared to young as analyzed by ELISA (Fig. 6F), consistent with increased autoantibodies and autoimmunity in aged mice.

Age-related partial denervation, signified by axonal blebbing at the neuromuscular junction (NMJ) followed by the partial loss of innervation, is a major contributor to muscle wasting. However, the pathobiology of motor neuron damage and cause of this denervation is unknown. We found that IgM accumulates in the motor neuron and fascia neighborhood of aged muscle (Fig. 6A). We therefore sought to determine whether IgM accumulation impacts age-related denervation. To test this, we performed immunostaining on whole mount EDL muscle isolated from young and aged mice, labeling acetylcholine receptors with bungarotoxin (BTX), the motor neuron with neurofilament, and IgM to assess whether IgM accumulates at the NMJ of aged muscle. We observed that while IgM staining is at background levels in young muscles, it is highly abundant in aged muscles (Fig. 6G-H). Indeed, our assay revealed a ∼4-fold increase in the mean fluorescence intensity of IgM at the NMJs of aged mice relative to young (Fig. 6H). To establish whether the IgM accumulation in aged muscles could affect motor neuron health, we stratified aged NMJs by the morphology of pre-synaptic neurofilaments at sites of axonal blebbing, which is indicative of axonal degeneration. Axonal blebbing at aged NMJs correlated with higher IgM signal. Since IgM can trigger the classical complement cascade (Chan et al., 2004; Daha et al., 2011; Sharp et al., 2019), we hypothesize that IgM accumulation underlies axonal degeneration in aged muscle. Together, our findings reveal an autoimmune origin for the pro-inflammatory changes commonly observed in aged muscle, likely mediated via IgM.

## Discussion

We present a spatial proteomic atlas of skeletal muscle regeneration at single cell resolution obtained using CODEX multiplex imaging and unbiased computational approaches (Fig. 1-2). Using a carefully curated panel of 32 antibodies, we reveal the spatial and temporal dynamics of 34 cell types during efficacious and dysregulated repair. By combining CODEX and new analytic tools, we show the power of this approach in resolving the spatial and temporal multicellular interactions in cellular neighborhoods that accompanies efficacious repair of muscle tissue damage and goes awry in muscle aging. Using neural network and unsupervised clustering, we are able to perform spatial pseudotime mapping of regeneration, creating a “regeneration clock” of cellular neighborhood interactions and how they change over time. These tools which enable an unbiased metric of the repair process, provide an integrated view of skeletal muscle tissue architecture at single cell resolution. Additionally, they serve as a resource for understanding complex changes in cellular neighborhoods in other tissues over time and during regeneration. Such information is lacking in widely used single cell approaches that entail tissue dissociation or non-multiplexed histological analysis. As such our atlas will serve as a reference for muscle biologists and a platform for discerning the biology underlying neuromuscular disease or regeneration in other tissues, and allow a holistic perspective of tissues that will inform therapeutic interventions.

Our atlas provides a spatial context for the heterogeneous cell populations that characterize muscle tissue, information that is lost upon dissociation of the tissue into single cells and nuclei for transcriptomic and proteomic studies (De Micheli et al., 2020a; Dos Santos et al., 2020; Giordani et al., 2019; Petrany et al., 2020a; Porpiglia et al., 2017). Knowledge of spatial relationships is critical to defining and characterizing specific cell-cell interactions in cellular neighborhoods. This is underscored by our ability to characterize localized biological regenerative processes, such as the clearance of ECM scaffolds and regeneration of muscle fibers (Fig. 3), the arrangement of cell types in dynamically changing cellular neighborhoods, and resolution of a temporal program of cell-cell interactions (Fig. 4D-E). By resolving the spatial temporal participation of cell types and functional subsets of cells, for the first time we can quantify localized regenerative activity (Fig. 4A-C) and identify tissue areas that deviate from the normal regeneration program (Fig. 4F-I). Our CODEX analysis of 34 cell types and characterization of 40 markers simultaneously in conjunction with the analytic tools we developed for analyzing this massive dataset enabled a previously unrecognized appreciation of the process of regeneration following muscle damage. Our data reveal the cellular mechanisms underlying impaired healing in the absence of macrophages in young (Arnold et al., 2007; Chazaud et al., 2003; Du et al., 2017; Ratnayake et al., 2021; Shang et al., 2020; Tidball and Welc, 2015; Tonkin et al., 2015). The impaired phagocytosis and accumulation of cellular debris that occurs after macrophage depletion fosters the dysregulated behavior that occurs across cell lineages (Fig. 3-5). We also pinpoint IgM accumulation as a novel feature of aging-associated chronic muscle inflammation that appears to be most prevalent at sites of denervation (Fig. 6).

By understanding and classifying the proximal enrichment of cell types during regeneration, our atlas reveals a specific temporal regenerative sequence of cell-cell interactions driven by directed signaling (i.e., signals from a source cell to a responder cell). Upon injury, injured myofibers recruit neutrophils, M1 macrophages, and FAPs and trigger endothelial expansion (Fig. 3B-C). The subsequent release of intracellular proteins, mitochondria, double stranded DNA and ATP create a damage associated molecular pattern (DAMP) recognized by innate immune cells and mesenchymal stem cells like FAPs (Vénéreau et al., 2015). Simultaneously, injured myofiber protein remnants acquire immunoglobulins (IgGs and IgMs) over time (Fig. 2B) and activate macrophages via FcR (Clynes et al., 1998; Deo et al., 1997). Neutrophils and macrophages release inflammatory lipid metabolites including prostaglandin E2 (PGE2) (Giannakis et al., 2019; Ho et al., 2017) and secrete cytokines (Arnold et al., 2007; Chazaud et al., 2003; Du et al., 2017; Ratnayake et al., 2021; Shang et al., 2020; Tidball and Welc, 2015; Tonkin et al., 2015) that activate MuSCs and promote their differentiation to proliferative myoblasts. Although cytokine signaling is necessary for MuSC expansion, the primary role of macrophages is to act as phagocytes that remove myofiber debris. Using clodronate liposomes to effect macrophage depletion, we find that IgM+ debris persists in injured myofibers, hindering the formation and maturation of new myofibers. Thus. our results demonstrate that phagocytosis is critical for clearing ECM scaffolds (Fig. 3E-H), a process required for their use as migratory tracts for myogenic progenitors to regenerate myofibers efficaciously (Webster et al., 2016).

Importantly, our data reveal for the first time that regeneration is not a series of checkpoints or extrinsic feedback driven process. Specifically, while the checkpoint and feedback models predict that disruption of regeneration would lead to a stall across cell types, we reveal that upon macrophage depletion, myogenic cells proceed through programmed differentiation and immune cell types mount an adaptive response (Fig. 4F-I). The lack of a stall leads to a co-existence of cell types that do not normally interact during regeneration. Macrophage depletion triggers the loss of coherent temporally and spatially coordinated cellular regenerative processes. Moreover, in aging we observe dysregulated changes to muscle structure and function that share features with young muscle that is in a persistent state of regeneration. Aged muscles exhibit greater myofiber turnover (Fig. 6A and 6H-I). They also feature an increased abundance of innate and adaptive immune cells, and altered ECM structures (Fig. S8A-B) consistent with an asynchronously regenerating tissue state (Fig. 6E-F). These changes are likely driven by combinations of systemic factors and local changes such as denervation, altered vascularization, and aberrant deposition of IgM.

Autoimmunity is common to a range of human neuromuscular disorders. For example, low muscle mass and sarcopenia develop in a significant proportion of rheumatoid arthritis patients, where IgM rheumatoid factor is a common diagnostic marker (Torii et al., 2019). Sjörgren’s syndrome, which can occur with rheumatoid arthritis, can also present with myositis with IgM expressing plasma cells infiltrating the muscle (Ringel et al., 1982). We found that IgM in the sera of aged mice exhibits rheumatoid factor activity (Fig. 6F), indicative of auto-reactivity. In our aged muscle data, the presence of IgM correlates with immune cell presence (Fig. 6C) and a chronic pro-inflammatory state. Additionally, the presence of IgM at the NMJ correlates with aberrant axonal blebbing (Fig. 6F,G) which could result from complement-mediated damage to the pre-synaptic cell membrane. Complement-mediated damage to neurons and muscle could cause to Ca2+ influx and negatively impact mitochondrial functions in the aged, which is a key therapeutic target for restoring neuromuscular function in the aged (Austin and St-Pierre, 2012; Palla et al., 2021). Taken together, our findings suggest that distal changes in the aged immune system could lead to age-related neuromuscular symptoms such as partial denervation and immune infiltration.

Taken together, our single cell resolution spatial atlas of skeletal muscle regeneration resolves temporally localized cell-cell dynamics. Combining CODEX multiplex imaging and neural network-powered computational approaches, we demonstrate that spatial analysis can reveal insights into dynamic processes involving multiple cell types and the tissue ECM in a manner previously not possible using approaches that dissociate tissues. These approaches pave the way for a better understanding of disease mechanisms, will improve diagnosis accuracy, and help validate drug effects across multiple cell types.

## Limitations of the study

In the current study, fluorescence intensity signals should not be interpreted as absolute quantification of protein expression. While most antibodies were validated to reflect their expected target, unexpected cross reactivity of a few antibodies was observed (e.g. anti-TNMD antibody showed high correlation with CD163 staining on M2 macrophages). This cross reactivity did not affect clustering or cell type designation, since our approach uses the expression of all markers, thus we were able to delineate M2 macrophages as CD163+ TNMD+ and tenocytes as CD163– TNMD+ (Fig. S4). Imaging artifacts and tissue folds were minimal but could affect the accuracy of quantification and cell type designation in the affected regions. Of note, signal for secreted proteins such as IgM, laminin, ERTR7 can localize to cells that do not express these proteins, which can explain discrepancies with transcriptome profiles of these cells. Although cell type annotations were manually validated with corresponding cell types in the tissue, algorithmic cell type identification is not perfect and remains an area of research and improvement.

## Methods

### Contact for reagent and resource sharing

Further information and requests for reagents and resources should be directed to and will be fulfilled by the lead contact, Helen M. Blau.

### Data and code availability

Spatial atlas of muscle cells and CODEX images have been deposited on Zenodo and are publicly available as of the date of publication. Due to the data size of raw CODEX datasets, down sampled versions of processed CODEX images are available as doi:10.5281/zenodo.6609234. Unprocessed or full resolution images are available from the lead contact upon request. Codes used for data analysis has been deposited at github.com/will-yx/. Any additional information required to reanalyze the data reported in this paper is available from the lead contact upon request.

### Experimental model and subject details

We performed all experiments and protocols in compliance with the institutional guidelines of Stanford University and Administrative Panel on Laboratory Animal Care (APLAC). Aged (24-28 mo.) mice C57BL/6 were obtained from the US National Institute on Aging (NIA) aged colony, and young (2-4 mo.) wild-type C57BL/6 mice from Jackson Laboratory.

### Experimental method details

#### Muscle injury and clodronate liposome injection

Muscle injuries were induced with a single intramuscular injection of 20 μL of notexin (5 μg/mL; Latoxan, catalog no. L8104) into the Tibialis anterior (TA) muscle. Injections were performed through the skin by inserting a 0.3 mL insulin syringe (BD; cat. 324702) from the distal point of the tibialis anterior (TA) muscle toward the knee, roughly parallel to the alignment of the myofibers. For macrophage depletion, 2 days after notexin injection, 40 μL of clodronate liposomes (Clophosome, FormuMax; cat. F70101C-N) or control liposomes (FormuMax; cat. F70101-N) was injected intramuscularly into the TA. Contralateral legs without injury or liposome injections were used as uninjured controls.

#### Construction of fresh frozen tissue section arrays

Muscle samples were dissected, embedded in a 15 x 15 mm tissue cassette filled with Tissue-Tek Optimal Cutting Temperature compound (VWR; 25608-930), and frozen in liquid nitrogen-cooled semi-frozen isopentane. Fresh frozen tissue samples were stored at -80 °C until processing. Tissue blocks were cryo-sectioned in a Leica CM3050S cryostat at 10µm thickness. Tissue sections were placed on square glass coverslips, 22 mm x 22 mm (Electron Microscopy Sciences; cat. 63757-10) pre-coated with poly-L-lysine (0.01% in ddH2O from 0.1% stock solution) mixture (Sigma; P8920). Single sections of a series of tissues of uninjured, day 1, 3, 6, and 10 post-injuries were arranged on each coverslip to form a tissue array. Each coverslip allowed for 4-6 tissues to fit, and they were stored at -20 °C until stained.

#### Traditional Immunofluorescence and antibody screening

Tissues were fixed with 4% PFA, blocked with blocking buffer (5% goat serum, 0.5% BSA, 0.5% Triton-X100 for 45 min in room temperature; if candidate antibody is a mouse IgG, 1% goat anti-mouse IgG Fab fragment (Jackson Research) was added. After blocking, tissues were washed with PBS (x3) and stained with candidate primary antibody for 2 hours at room temperature or overnight at 4 °C, washed with PBS and stained with appropriate secondary antibody for 2 hours in room temperature. DAPI and TrueBlack stain were added, and then the slides were mounted and inspected under the microscope. This was done to determine whether the antibody stained the relevant target and to decide on the dilution ratio for the CODEX staining. Prior to adding the antibodies to the CODEX antibody panel, all antibodies were tested on mouse skeletal muscle sections for their staining efficiency following an IHC staining protocol, as follows.

#### CODEX Buffers and solutions

Buffers and solutions were prepared as described in Schürch et al. 2020. All buffers were filtered sterile using 500 mL 0.2 μm pore size filters and stored at room temperature unless otherwise specified.

Hydration Buffer (S1), Staining Buffer (S2), and Storage Buffer (S4); (Akoya Biosciences).

TE buffer: 10 mM Tris pH 8.0, 1 mM EDTA in ddH_2_O (Invitrogen).

CODEX buffer (H2): 150 mM NaCl, 10 mM Tris pH 7.5 (Teknova), 10 mM MgCl_2_ · 6 H_2_O (Sigma), 0.1% w/v Triton X-100 (Sigma) and 0.02% w/v NaN_3_ in ddH_2_O; stored as a 10x stock solution.

Blocking component 1 (B1): 1 mg/ml mouse IgG (Sigma) in S2.

Blocking component 2 (B2): 1 mg/ml rat IgG (Sigma) in S2.

Blocking component 3 (B3): Sheared salmon sperm DNA, 10 mg/ml in H_2_O (Thermo Fisher).

Blocking component 4 (B4): Mixture of non-modified CODEX oligonucleotides at a final concentration of 0.5 mM in TE buffer (IDT).

CODEX plate buffer: 33.3 nM Hoechst 33342 (Thermo Fisher) and 0.5 mg/mL B3 in 1x CODEX buffer.

F fixative solution (BS3): 200 mg/ml BS3 (bis(sulfosuccinimidyl)suberate; Thermo Fisher) in anhydrous DMSO (Sigma); stored at -20°C in 15 μL aliquots; used freshly thawed.

CODEX antibody stabilizer solution: Antibody Stabilizer in PBS (Candor Bioscience) with 5mM EDTA and 0.01% sodium azide (Sigma).

#### Generation of CODEX DNA-conjugated antibodies

Antibody conjugations were performed, and oligonucleotide barcode sequences were as described in Schürch et al. 2020. Oligonucleotide barcodes were conjugated to antibodies via maleimide-thiol reactions.

Protected 5’ maleimide-modified oligonucleotides were purchased from Trilink or GeneLink and were deprotected according to manufacturer’s protocol. In brief, lyophilized oligonucleotides were washed in anhydrous acetonitrile, then heated to >90 °C in anhydrous toluene for 4h (with an exchange with fresh toluene after 2h). Deprotected oligonucleotides were washed in anhydrous ethanol three times, resuspended in Buffer C (150 mM NaCl, 2 mM Tris (from a 50 mM stock solution, pH 7.2), 1 mM EDTA and 0.02% w/v NaN_3_ in ddH_2_O), aliquoted to PCR strip tubes (100 μg per aliquot), then snap frozen by liquid nitrogen and lyophilized overnight on a FreeZone 4.5 Plus lyophilizer (Labconco). Lyophilized deprotected oligonucleotides were stored at -20°C until antibody conjugation.

50 or 100 μg of a validated antibody was placed in an Amicon Ultra 0.5 mL 50 kDa molecular weight cutoff (MWCO) spin column (EMD Millipore; cat. UFC505096) and concentrated by centrifugation at 12000 g for 8 min. Antibodies with BSA or glycerol contaminants were first concentrated in a 100 kDa MWCO spin column (EMD Millipore; cat. UFC510096) and washed twice with 400 μL of PBS before being transferred to the 50 kDa column. MWCO filters were first conditioned with 500 μL of PBS-tween and spun down at 12000 g for 2 min. All centrifugation steps were at 12,000 x g for 8 min and flow-through was discarded, unless otherwise specified. To reduce disulfide bonds to free thiols, a mixture of 12.5 mM TCEP and 2.5 mM EDTA in 1X PBS was added to the concentrated antibody on the spin column and incubated for exactly 30 min. Columns were centrifuged to remove the TCEP and washed with buffer C (150 mM NaCl, 2 mM Tris stock solution, pH 7.2, 1 mM EDTA and 0.02% w/v NaN_3_ in ddH_2_O). Per 50 μg of starting antibody, 100 μg of lyophilized deprotected maleimide oligonucleotides was reconstituted in 15 µl UltraPure Distilled Water (Invitrogen) and then mixed with 330 µL Buffer C and 50 µL 5M NaCl. The oligonucleotide mixture was added to the reduced antibody and incubated at room temperature for 2 h. The conjugated antibody was spun down and washed three times with 450 µl High salt PBS (PBS with 1M NaCl). Per 50 µg of starting antibody, an amount of 100 µl of CODEX antibody stabilizer solution was added to the column, mixed by pipetting, then inverted into new collection tubes and centrifuged at 4,000 x g for 2 min. Conjugated antibodies were stored at 4 °C. Antibody-oligonucleotide barcode combinations are listed in Table S1.

#### Tissue processing and staining for CODEX

Antibody staining was performed in two staining steps. Antibody Mix 1 (AM1) was prepared by pipetting Myod1, MyoG, DMD, eMyHC, p-H3, Itga7, Pax7, PDGF-alpha, Laminin a2, MyHC antibodies at indicated dilutions in Table S1 in S1 buffer and mixed with blocking reagents (B1, B2, B3, B4) in a ratio of 210:10:10:10:10. Antibody Mix 2 (AM2) was prepared by mixing B220, CD11b, CD3, CD4, CD8a, ERTR7, CD29, CD11c, CD16/32, IgM, MHCII, Ter119, CD38, GFAP, F4/80, Ki67, Ly6G, Sca1, CD45, CD90, CD47, CD31, CD163, CD9 antibodies at indicated dilutions in S2 buffer and blocking reagents.

Tissue section arrays were thawed for 2 min and washed twice with Hydration Buffer (S1). Sections were fixed with 1.6% PFA in S1 for 10 min, then washed with S1. 150μL of AM1 was added to each coverslip and incubated in a hydration chamber at 4 °C overnight. Coverslips were washed with S1 and then S2 buffer. 150 μL of AM2 was added to each coverslip and incubated in a hydration chamber at room temperature for 3 h. Coverslips were then washed twice in S2, fixed with 1.6% PFA for 10 min, and washed 3 times with PBS. Tissues were then fixed with ice cold methanol for 5 min, followed by PBS washing. To reduce autofluorescence, 200 μL of TrueBlack (Biotium) was added to the coverslips for 1 min according to manufacturer’s recommendations. A final fixation step was performed F Fixative for 20 min followed with PBS washing. Coverslips were stored at 4 °C in Storage Solution (S4) until imaging.

#### CODEX multi-cycle reaction and image acquisition

CODEX multi-cycle reactions were performed on an Akoya Bioscience CODEX instrument, according to manufacturer’s instructions, and imaged on an automated Keyence BX-700 microscope equipped with a Nikon 20x NA 0.75 Plan APO len. Cycle arrangement and reporter plate setups are described in Table S2. 10 μL of each corresponding fluorophore-conjugated oligonucleotide reporter to antibodies (10 μM in TE buffer; IDT) of a given cycle was mixed with CODEX plate buffer to a total volume of 250 μL and arranged in a 96-well round bottom plate. The first cycle, and second and third last cycles designated as blank cycles to capture autofluorescence, no reporter oligonucleotides were added. The final cycle, 1 μL of DRAQ5 was added to 249 μL of CODEX plate buffer.

Fluidics exchange and image acquisition were fully automated through the Akoya Biosciences CIM software and Keyence Microscope BZ-X Viewer software. Each tissue was imaged in a 5x7 tiled region and with 33 z-slices with an axial resolution of 0.8 μm. Imaging regions were manually selected after initial staining with Hoechst to capture as much of each tissue as possible. The z-position of the tissue was automatically determined by the autofocus feature on the Keyence software, on the center tile of each imaging region, every cycle.

#### Immunofluorescence of muscle fiber bundles

Extensor digitorum longus (EDL) muscles were dissected and fixed in 4% PFA in PBS for 20 min, then washed with PBS. EDL muscles were manually teased into myofiber bundles under a dissection microscope avoiding any contact with the endplate band. Muscle bundles were permeabilized in 0.3% Triton X-100 in PBS (PBS-T) for 30 minutes and nonspecific binding was blocked using goat serum-based blocking solution (5% goat serum in PBS-T) for 1 hour. Tissues were incubated with antibodies against neurofilament (2H3, DSHB, mouse IgG1) and synaptic vesicle (SV2, DSHB, mouse IgG1) in blocking solution at 5 ug/ml for minimum of 24 hours at 4C on a rocker. Muscle fiber bundles were washed extensively with PBS-T and stained in suspension with Alexa Fluor 546 conjugated goat anti-mouse IgM and Alexa Fluor 488 conjugated goat anti-mouse IgG subclass1 antibodies and Alexa Fluor 647 conjugated bungarotoxin (BTX) in blocking solution. After extensive washing in PBS muscle fiber bundles were mounted onto SuperFrost Plus slides (Fisher, cat. 12-550-15) using Fluoromount G (Thermo Fisher; cat 00-4958-02). Muscles were imaged on a spinning disc confocal microscope. Z stacked 3D images were processed by maximum intensity projection. Neuromuscular junctions (NMJ) were masked by thresholding on BTX staining and intensities for IgM was measured in ImageJ. NMJ fragmentation, axonal blebbing phenotype were manually scored.

#### ELISA for IgM Rheumatoid Factor

Blood from young and aged mice were collected by cardiac puncture in non-heparin tubes, allowed to clot for 30mins, and spun down at 3000 x g for 10 min. Sera was collected as the supernatant, snap frozen, and stored at -80°C until analysis. On the day of analysis, sera were thawed on ice and IgM Rheumatoid Factor Mouse ELISA (BioVendor; cat 634-02689) was performed according to manufacturer’s instructions.

### Quantification and Statistical Analysis

#### Computational image processing and in silico autofluorescence clearing

CODEX images from repeated imaging cycles were processed using the CRISP Image Processor as described in Palla et al., 2021. Hoechst channels from each imaging cycle was used to align tiles within each tissue region, 3D drift compensation across cycles, and identify slice(s) of best focus in the Z axis. All registration and alignment steps were performed in Fourier transformed frequency domain at sub-pixel resolution. Each image stack was then deconvolved using Richardson-Lucy algorithm over 50 iterations with a computed vector point spread function (PSF) estimated using a Gibbson-Lanni model with de-ringing filters. Gibson-Lanni parameters were estimated based on the imaging conditions (xy-resolution of 0.37744 μm per pixel, z-resolution of 0.8 μm per slice, working distance (ti0) of 1000 μm, relative position of the tissue (zpos) of 10 μm, coverslip thickness (tg) of 170 μm, glass refractive index (ng) of 1.500, immersion refractive index (ni) of air 1.0003, and sample refractive index (ns) of CODEX buffer containing 20% DMSO ∼1.397). PSFs were generated with 1000 Bessel functions (nbasis) and 1000 computed angles (nrho). Independent PSFs were generated for each channel according to the emission wavelengths of the fluorophore and their full-width half-max emission as follow: Hoechst, 455 ±70 nm; FITC, 517 ±40 nm; ATTO550 or Cy3, 580 ±30 nm; Alexa647, 675 ±25 nm. During deconvolution, images were translated in Z axis in the frequency domain to fit the best focus slice for the entire tissue to a single plane. Concurrently, images were corrected for lens and microscope sensor artifacts using pre-generated flatfield and darkfield images, respectively. After deconvolution, images were re-registered in the X-Y axes, then registered across all channels and stitched into full resolution mosaics. After stitching, blank cycles imaged at the beginning and end cycles were used to subtract autofluorescence for each channel of each cycle. Linear interpolation of imaging time and exposure time was used estimate the autofluorescence contribution of signal and this signal was subtracted from each channel at single pixel resolution.

#### Neural network identification of nuclei and tissue features

Nuclei were segmented using a modified version of CellSeg as described in Lee et al., 2021. The segmentation portion of CellSeg was run using pre-trained models on the full resolution CRISP stitched image of the DRAQ5 channel with the following parameters: overlap of 80, min_area of 40, increase_factor of 3, autoboost_percentile of 99.98.

Tissue features were classified from select channels of the CODEX dataset using FiberNet. FiberNet is a neural network classifier that performs semantic segmentation on multi-channel images of stained tissue sections. FiberNet was trained to identify healthy, regenerating, injured, and ghost muscle fibers along with stroma, motor neurons, and background areas of tissues.

The FiberNet model architecture is based on a residual neural network (ResNet) with modifications to improve rotational invariance. The network determines an object class for each pixel in the 32-channel input image, along with a confidence score for each of the eight possible classes. For each image position an input window samples the source image stack at 1 and 1/4 scale, which gives the model access to local detail as well as broad context of the surrounding area. Input data is fed to two parallel paths within the network, the first of which splits the image into four quadrants and rotates them to enforce rotational invariance. Because the main branch only sees a quarter of the input window, a supplemental branch allows the network to consider the entire breadth of the input field of view, albeit at a lower resolution. The primary ’quadrant’ branch employs a series of 2D convolutions with shortcut connections typical of a ResNet architecture. This branch then bifurcates, processing the data as well as its transpose through a final 2D convolution layer and two dense layers. The outputs of the four quadrant branches were stacked, and the mean, minimum, and maximum values were computed across the eight quadrant results. This again enforces rotational invariance between quadrants. The ’overview’ branch computes a row-wise and column-wise mean of the data and concatenates the left and right and top and bottom halves of these averages into a single tensor. Once again, only the mean, minimum, and maximum values of these four sections were taken to enforce rotational invariance. The network then concatenates the dense output of the two data paths and continues through a further series of dense layers and the final categorical output layer. The exact dimensions of the model were parametrized based on the number of convolution layers, method of padding used, and size of the ResNet output. A typical input window size is 85x85 pixels with 1024 channels at the output of the convolution layers. FiberNet was trained on expert curated CODEX data on NVIDIA 2080Ti GPUs using Tensorflow (https://www.tensorflow.org). Interpretation was performed on full resolution multi-channel CODEX images to classify each pixel in the image. Results were manually reviewed and validated by experts to assure accuracy. A FiberNet Lite model was trained using only the Laminin, DNA, and autofluorescence channel to allow segmentation of myofibers in traditional immunofluorescence images of skeletal muscle cross sections. Neural network classified image were post-processed into morphological masks, refined using morphological erosion and dilation functions from the scikit-image package, and morphologically assessed for area, centroid, mean intensity, major and minor axis lengths using the measure.regionprops_table functions from the Scikit-Image (Walt et al., 2014) python package. Mask objects were filtered by area for greater than 1000 and less than 30000 pixels.

#### Quantification of antibody staining intensity

Nuclei masks segmented by CellSeg were used to quantify antibody staining intensity. Nuclei masks were expanded using morphological growth by 2.5 pixels. The border 2 pixels of the grown mask were then used to quantify cytoplasmic or membrane staining intensity of each antibody, and the remaining interior pixels were used to quantify nuclear staining of each antibody. The mean value of pixels in each compartment for each cell was used for downstream analysis.

#### Data preprocessing, unsupervised clustering, and annotation of single cells

Single cell staining intensity data across 11 CODEX experiments (“run”) consisting of 47 tissues including young uninjured, young regenerating, young regenerating after clodronate treatment, and aged uninjured samples were concatenated together and analyzed as a single dataset using HFcluster. Highly autofluorescent cells and falsely segmented cells were removed based on intensities measured in the blank channels and lack of signal in DNA channels. Staining intensities for single cells in each run were quantile normalized to 95th percentile and zero centered at the median or 50^th^ percentile. Normalized intensities of select markers (Table S1) were used for clustering. For HFcluster’s two step clustering approach, an automated elbow-finding approach is used to estimate a threshold for high confidence positive staining. Cell intensities were high pass filtered at the threshold value. This step sparsifies the intensity matrix causing cells with low or poor staining patterns to drop out. The filtered intensity values were clustered with the Louvain algorithm (as implemented in the single cell analysis python package, Scanpy (Wolf et al., 2018)), whereby poorly stained cells will cluster together and cells with high confidence staining can be more accurately clustered. The cluster labels of cells with high confidence staining were propagated onto poorly stained cells using the pre-filtered normalized intensity matrix. The propagated clusters were merged via hierarchical clustering using correlation distances of the mean intensities of each cluster, which resulted in 75 clusters. Merged clusters were then manually annotated based on expected antibody staining patterns into 33 cell subsets. Each subset was validated against the original CODEX image data for accuracy of annotation.

#### Generation of cell-cell interaction networks and cellular neighborhoods

Cell-cell interaction networks were predicted based on the spatial arrangement of cell types within tissues. For each cell in the dataset, a niche or window consisting of the index cell and its 10 nearest neighbors were identified using a distance map of cells within its source tissue. The cell type identities of the neighbors were counted to reveal the niche composition of each cell. To normalize for differences in abundances of each cell type, the niche composition was quantile normalized and the enrichment of pairwise interactions were shown as mean quantile values for niches of all cells of a given cell type.

Cell neighborhoods were defined by clustering niches according to cell type compositions as described in Schürch et al. 2020 using modified clustering approaches. In brief, niches for all cells were clustered using the Leiden algorithm with resolution of 0.5. Scarce neighborhoods with a total of less than 1000 cells across all tissues were merged into an unassigned cluster. The abundance of neighborhoods in each tissue across the regeneration time course was used to meta-cluster neighborhoods into temporal clusters.

#### Tissue cell type composition and enrichment analysis

Cell types and numbers of cells in each cell neighborhood were counted in each tissue, normalized to the total number of cells in the tissue, which represents the proportion of cells in the tissue. Log transformed enrichment for any given cell type or cell cluster across time is calculated as:

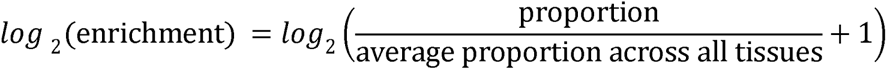

#### Spatial pseudotime analysis

Pseudotime encoding for each cell type was calculated as a likelihood of cell type at each sampled time point. To normalize for the different numbers of cells at given time points, cell type counts were first normalized to the total number of cells found at a given time point. Then, the likelihood of a cell type being present in samples of a given time point was calculated as normalized counts divided by the sum of normalized counts across the entire normal time course of regeneration. The average pseudotime of a cell type was calculated by multiplying the likelihood with regeneration time points (days after injury; an estimated value of day 20 was used for uninjured muscle). For tissue pseudotime, the positional information of each cell in the tissue is encoded with their average pseudotime according to their cell type. Tissues were then subsampled into ∼75 x 75 μm (200 x 200 pixel) bins, whereby the mean pseudotime of cells within each bin, difference of the mean pseudotime to actual time, and variance of pseudotimes of cells within each bin were assessed using Numpy (Harris et al., 2020) and visualized using the Scikit-Image (Walt et al., 2014) python package. Bins lacking cells, thus resulting in NaN values, were ignored in the subsequent analysis.

#### Dimensional reduction using UMAP and clustering for co-occurring cell types within neighborhoods and ECM scaffolds

Dimension reduction for the cell type compositions of tissues and cells within ECM scaffolds was performed using Uniform Manifold Approximation and Projection (UMAP) (McInnes et al., 2020). Input data were normalized along dimensions as proportions of total events or as enrichment across time points. Minimum distance and number of neighbors parameters were adjusted according to number of data points to maximally resolve heterogeneity. Clustering was performed on the UMAP embedding using the Leiden algorithms.

#### Statistical analysis

Statistical analysis was performed with one-way ANOVA with multiple pairwise comparisons Tukey HSD tests using the Scipy stats module (https://scipy.org) and bioinfokit (Bedre, 2021) python packages. P values of less than 0.05 were considered statistically significant.

## Acknowledgements

We apologize to those investigators whose important work we were unable to cite or describe owing to space constraints. We thank Akshay Balsubramani for their critical input and advice on computational approaches to resolve single cell data. We thank Megan Mayerle for careful editing and proofreading, Peggy Kraft and Kassie Koleckar for technical support, and Cindy Paulazzo and Megan Mayerle for administrative support. This study was supported by the Baxter Foundation, the Li Ka Shing Foundation, Milky Way Research Foundation MWRF-216064, California Institute for Regenerative Medicine (CIRM) grant DISC2-10604 and US National Institutes of Health (NIH) grant 5R01AG020961, 1R01AG069858 to H.M.B and 5R01HG009674 to A.K. and H.M.B. and U.S. Food and Drug Administration (FDA) Medical Countermeasures Initiative contracts 75F40120C00176 and HHSF223201610018C to GPN. Y.X.W. was supported by the Canadian Institutes of Health Research, Stanford Translational Research and Applied Medicine (TRAM) Pilot grant, and NIH K99 award K99NS120278, J.G was supported through the DARE Fellowship Program, a partnership between the Lundbeck Foundation and Innovation Centre Denmark, based in Silicon Valley. C.M.S. was supported by an Advanced Postdoc Mobility Fellowship from the Swiss National Science Foundation (P300PB_171189, P400PM_183915). Y.G. was supported by NIH grants U54 CA209971, 5U01AI101984 and 3U54HG010426 to G.P.N. This article reflects the views of the authors and should not be construed as representing the views or policies of the FDA, NIH or other funding sources listed here.

## Author Contributions

Y.X.W. and H.M.B. conceived and designed the study. Y.X.W., J.N.H., C.M.S., and Y.G. developed assays and optimized conditions for CODEX. Y.X.W., J.N.H, J.G., carried out muscle regeneration studies and performed CODEX imaging. Y.X.W., and C.H. developed CRISP, FiberNet, and HFCluster and carried out computational analysis of CODEX data. M.Y.L., C.M.S., Y.G., and G.P.N. developed CellSeg, which was modified by C.H. for this study. Y.X.W., M.A., and S.S. performed analysis of aged muscles and performed analyses of neuromuscular junctions. Y.X.W., J.G., C.H., and H.M.B. wrote the manuscript with inputs from all authors.

