## Supplemental Information for "A single cell spatial temporal atlas of skeletal muscle reveals cellular neighborhoods that orchestrate regeneration and become disrupted in aging"

**Supplemental Information and Data**

Supplemental Figures S1-S7

Supplemental Tables:
Table S1. Primary antibodies used for CODEX.
Table S2. CODEX multicycle setup.

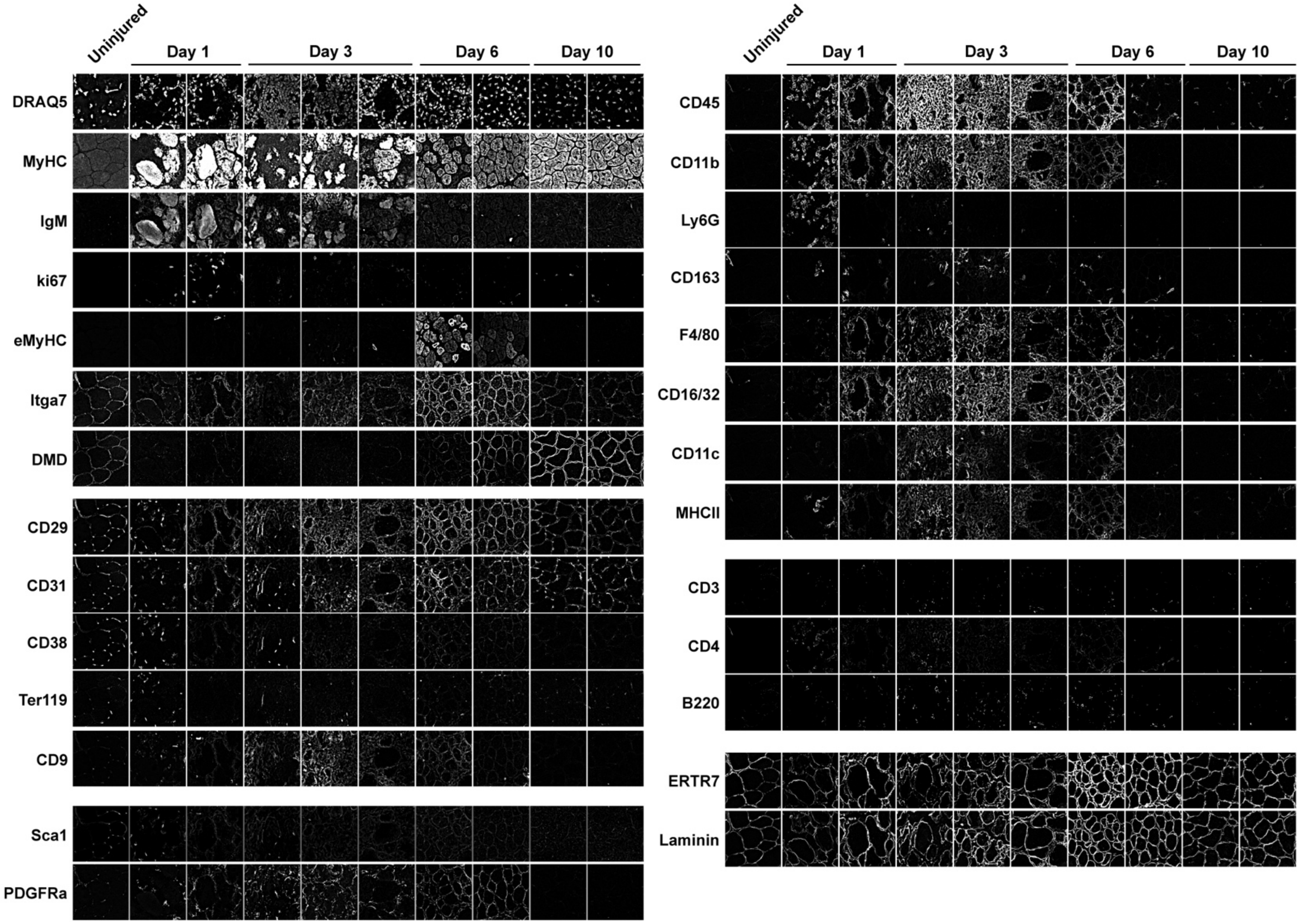


**Supplemental Figure S1. Temporal dynamics of markers of skeletal muscle cell types across regeneration time points resolved by CODEX.** Related to Figures 1 and 2.

Representative CODEX images of uninjured and regenerating muscle after 1, 3, 6, and 10 days after injury. The same field-of-view is shown in each column.

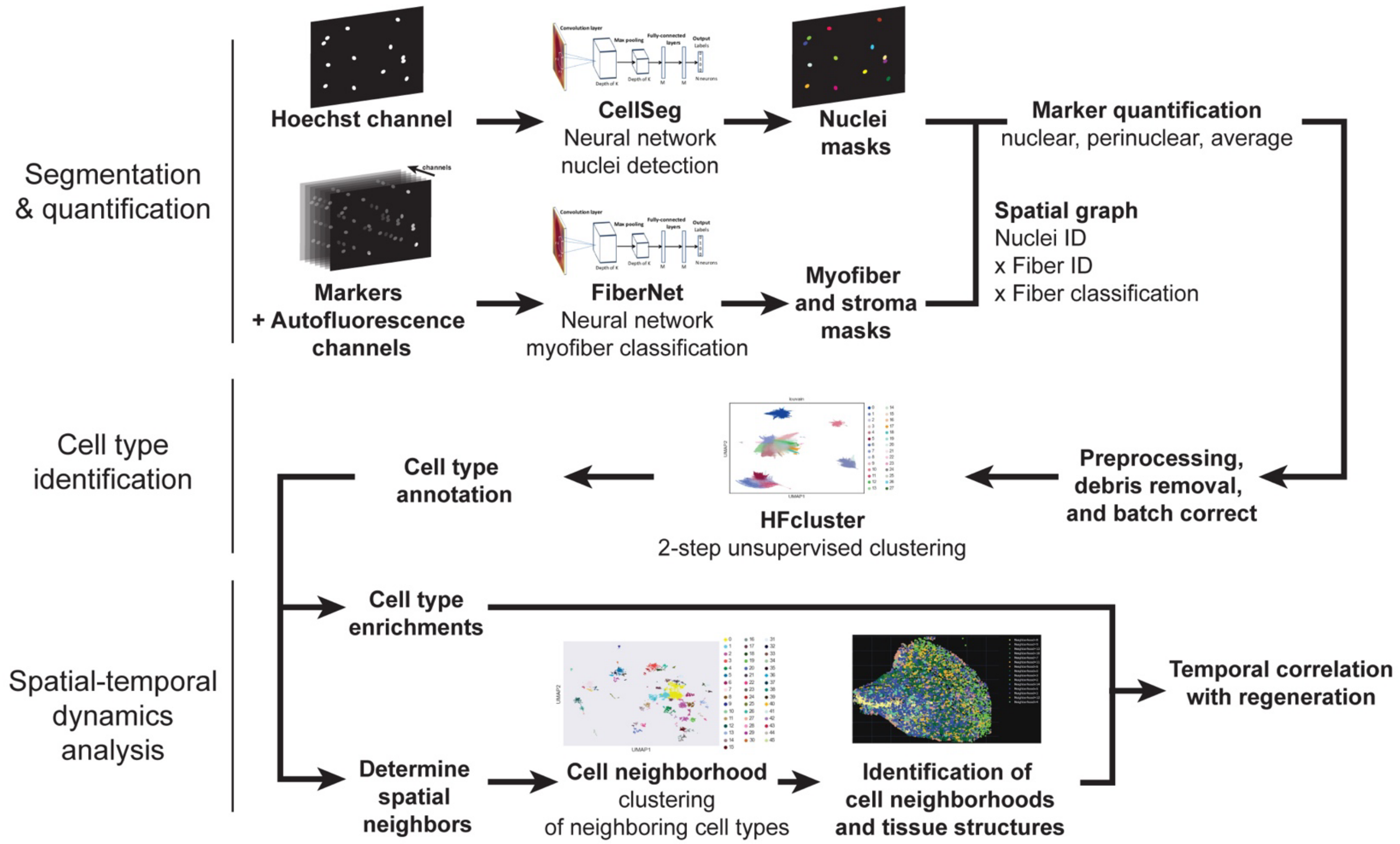


**Supplemental Figure S2. Computational pipeline to segment, quantify, identify, and annotate cell types found in skeletal muscle regeneration and resolve intercellular spatial temporal relationships.** Related to Figure 2.

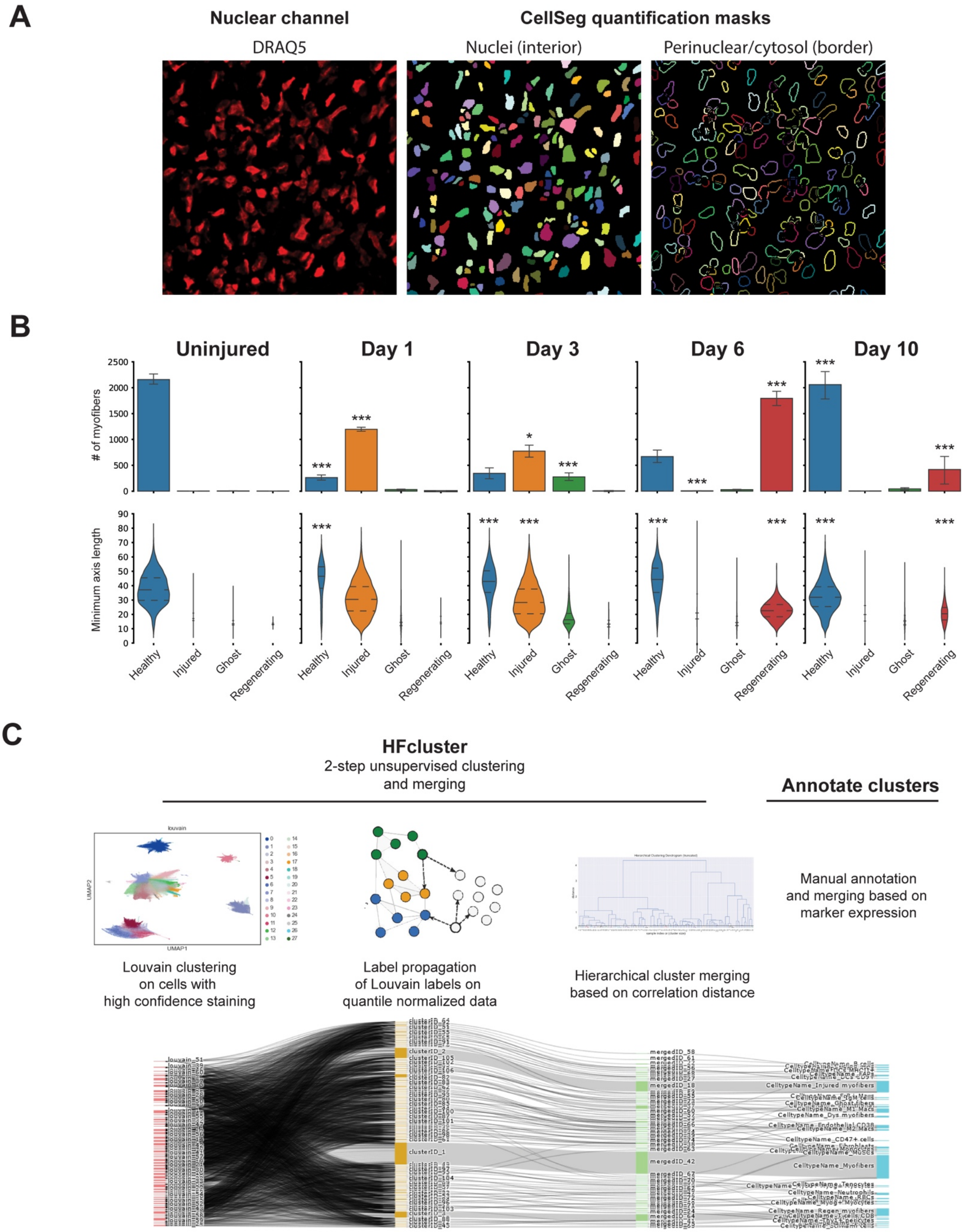

A
Nuclear channel
CellSeg quantification masks
DRAQ5
Nuclei (interior)
Perinuclear/cytosol (border)
B
Uninjured
Day 1
Day 3
Day 6
Day 10
### of myofibers
2500
2000
1500
1000
500
0
***
*
Minimum axis length
90
80
70
60
50
40
30
20
10
0
Healthy
Injured
Ghost
Regenerating
C
HFcluster
2-step unsupervised clustering
and merging
Annotate clusters
louvain
UMAP1
UMAP2
Louvain clustering
on cells with
high confidence staining
Label propagation
of Louvain labels on
quantile normalized data
Hierarchical cluster merging
based on correlation distance
Manual annotation
and merging based on
marker expression
clusterID_2
clusterID_1
mergedID_58
mergedID_18
mergedID_42
CelltypeName_Injured myofibers
CelltypeName_CD47+ cells
CelltypeName_Myofibers

**Supplemental Figure S3. Algorithmic segmentation of nuclei and cells, and cell type annotation.** Related to Figure 2.

**A)** Representative performance of CellSeg in segmenting nuclei from CODEX images based on DRAQ5 nuclear staining (left). Nucleai (middle) and perinuclear/cytosol (right) masks used to quantify antibody staining.

**B)** Quantification of myofiber classes identified by FiberNet in uninjured and regenerating muscles at day 1, 3, 6, and 10 after injury (top). Minimum axis lengths of myofiber classes identified by FiberNet in uninjured and regenerating muscles at day 1, 3, 6, and 10 after injury (bottom). Statistics indicate significant change from the previous time point; n=4-8 per group; * p<0.05; ** p<0.01; *** p<0.005.

**C)** Processes used for HFCluster and cluster merging and Sankey diagram of cell labels through the clustering steps.

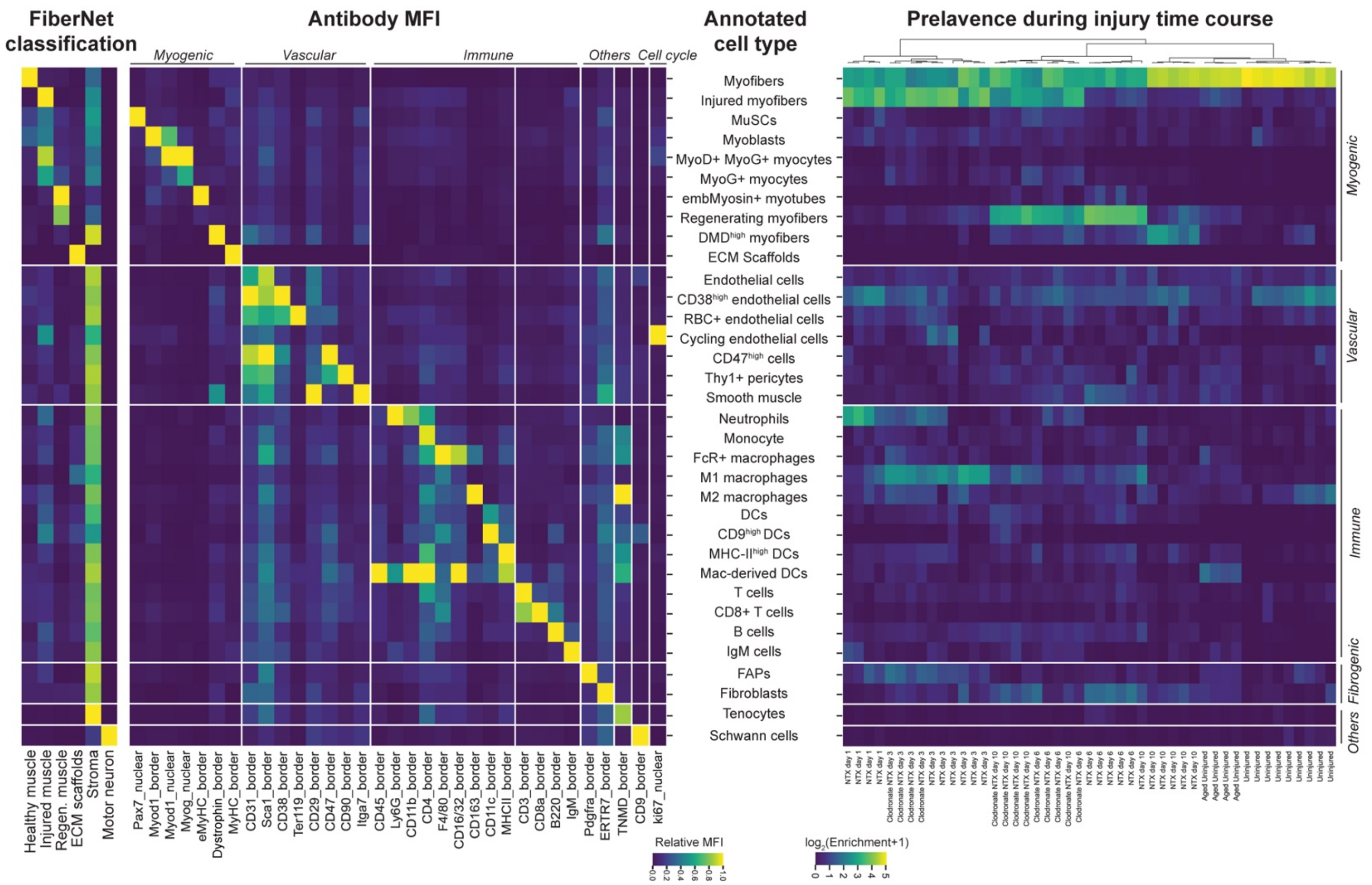


**Supplemental Figure S4. FiberNet classification, marker expression, and relative abundance of cell types found in uninjured, regenerating, and aged skeletal muscles.** Related to Figure 2.

Heatmap of mean FiberNet classification (left) and antibody staining intensities (middle left) for each annotated cell type resulting from HFCluster. The relative abundance of each cell type in each tissue is shown in a heatmap on the right. The cellular composition of the tissues cluster by the time point, regeneration progress, age, and clodronate treatment. n=4-8 per group.

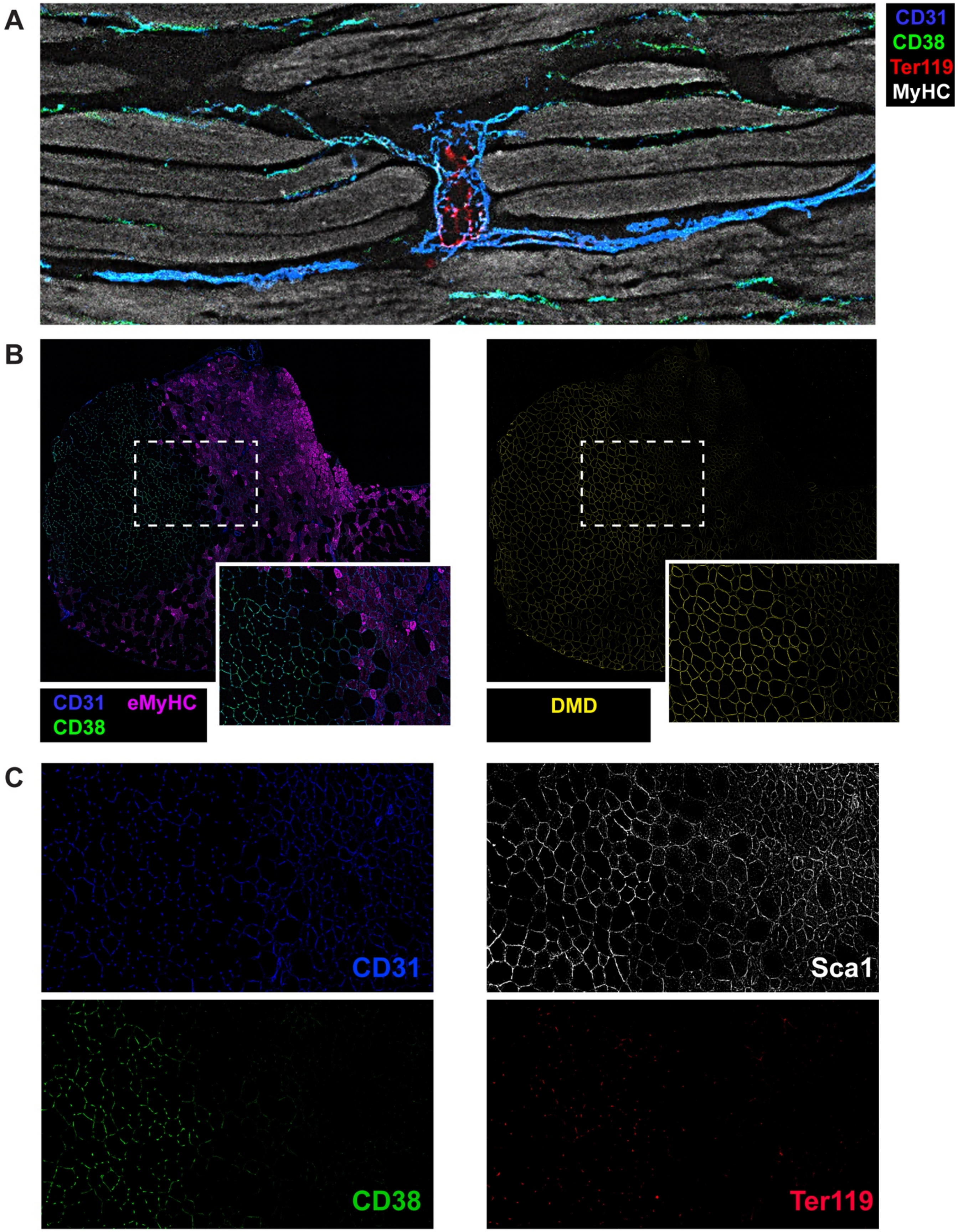
A
CD31
CD38
Ter119
MyHC
B
CD31
eMyHC
CD38
DMD
C
CD31
Sca1
CD38
Ter119

**Supplemental Figure S5. Identification of CD38 as a marker of perfused capillary endothelial cells found in skeletal muscle.** Related to Figure 2.

**A)** Representative CODEX image of longitudinally sectioned uninjured muscle showing CD31+ (blue) CD38+ (green) capillaries connecting to larger CD31+ CD38– vessels containing Ter119+ (red) red blood cells (RBC). Myosin heavy chain (MyHC) in gray.

**B)** Representative CODEX image of regenerating muscle day 6 after injury. Injured and regenerating regions of the muscle are marked by embryonic forms of myosin heavy chains (eMyHC; magenta, left panel) which anti-correlates with CD31+ (blue) CD38+ (green) capillaries and dystrophin staining (yellow, right panel). Inset is an enlarged view of the region within the dashed box, also shown in C.

**C)** Enlarged images of endothelial markers of the inset from B. Comparable staining for endothelial cell markers CD31 (blue) and Sca1 (grays) can be found across the boundary of injury. CD38 (green) and Ter119 (red) colocalize in capillary endothelial cells only on the uninjured regions.

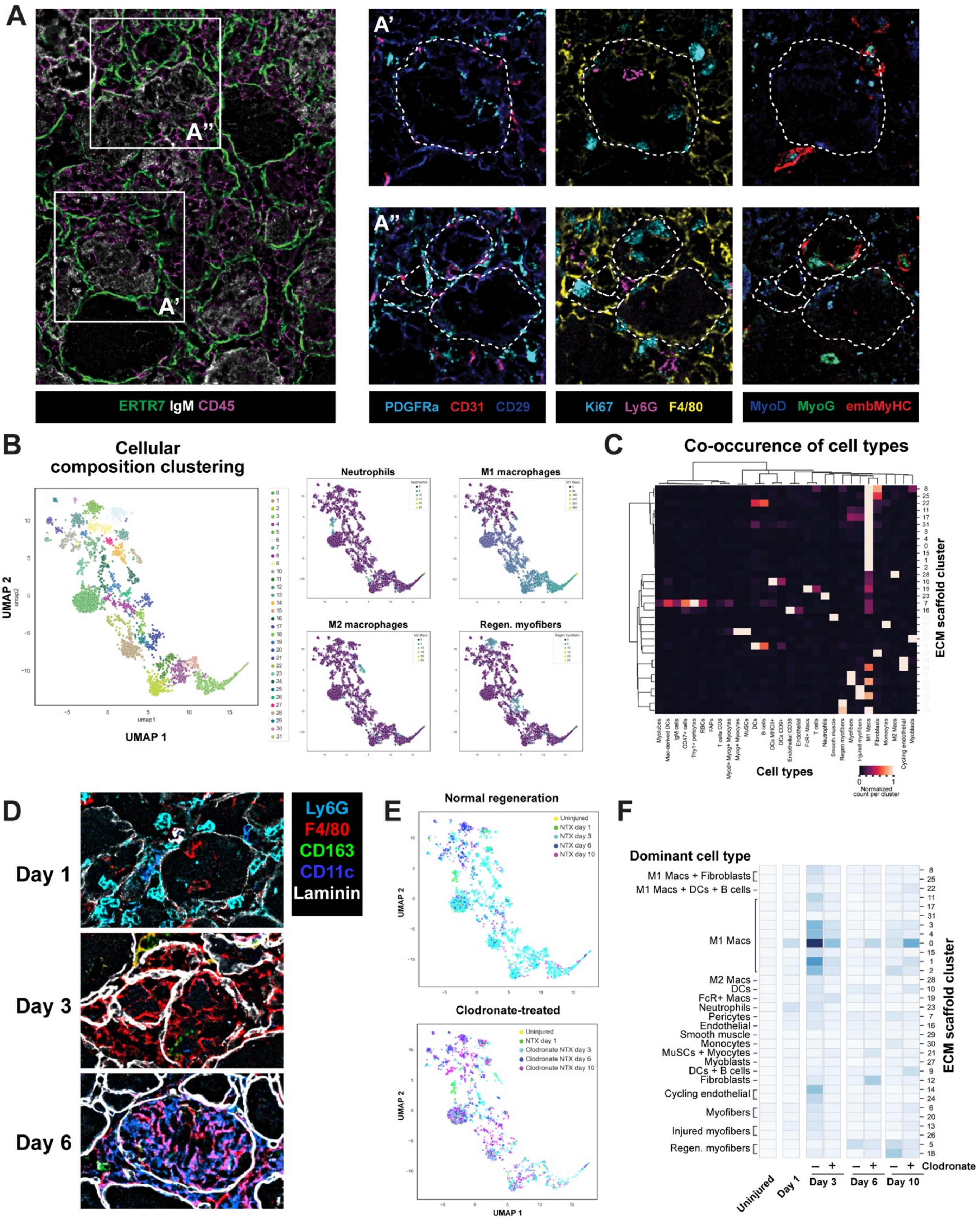
A
A''
A'
ERTR7 IgM CD45
PDGFRa CD31 CD29
Ki67 Ly6G F4/80
MyoD MyoG embMyHC
B
Cellular composition clustering
UMAP 1
UMAP 2
Neutrophils
M1 macrophages
M2 macrophages
Regen. myofibers
C
Co-occurence of cell types
Cell types
ECM scaffold cluster
Normalized count per cluster
D
Day 1
Day 3
Day 6
Ly6G
F4/80
CD163
CD11c
Laminin
E
Normal regeneration
Uninjured
NTX day 1
NTX day 3
NTX day 6
NTX day 10
Clodronate-treated
Clodronate NTX day 3
Clodronate NTX day 6
Clodronate NTX day 10
F
Dominant cell type
M1 Macs + Fibroblasts
M1 Macs + DCs + B cells
M1 Macs
M2 Macs
DCs
FcR+ Macs
Neutrophils
Pericytes
Endothelial
Smooth muscle
Monocytes
MuSCs + Myocytes
Myoblasts
DCs + B cells
Fibroblasts
Cycling endothelial
Myofibers
Injured myofibers
Regen. myofibers
Clodronate
Uninjured
Day 1
Day 3
Day 6
Day 10
ECM scaffold cluster

**Supplemental Figure S6. Dynamics of cells traversing the extracellular scaffold of injured myofibers.** Related to Figures 3 and 4.

**A)** Representative CODEX image of ECM scaffolds in regenerating muscle day 3 after injury. ECM marked by ERTR7 (green); injured myofibers marked by IgM (grays); and immune cells marked by CD45 (magenta). Regions in the outlined boxes are shown in **A'** and **A''**.

**A' and A'')** Enlarged views of boxed regions from panel **A** showing a range of cell type traversing the ECM scaffold. PDGFRa+ FAPs (cyan, left); CD31+ ECs (red, left); Ly6G+ neutrophils (magenta, middle); F4/80+ macrophages (yellow, middle); MyoD+ MyoG+ myogenic progenitors (blue and green, right); and eMyHC+ myotubes (red, right). Ki67 marks dividing cells. Dashed outlines are drawn based on ERTR7 staining from panel **A**. The same field-of-view is shown in each row.

**B)** UMAP embedding and clustering for the heterogeneity of ECM scaffold based on cell compositions within them. Each dot is one ECM scaffold containing a range of cells as shown in panel **A**. Clustering identified 31 types of ECM scaffolds with varying cell type compositions (left). UMAP embedding of ECM scaffolds containing neutrophils, M1 macrophages, M2 macrophages, and regenerating myofibers shown on the right.

**C)** Heatmap showing the co-occurrence of cell types within each ECM scaffold cluster from panel **B**.

**D)** Representative CODEX image of myeloid cell types traversing ECM scaffolds in regenerating muscle day 1, 3, and 6 after injury. ECM marked by laminin (grays); neutrophils marked by Ly6G (cyan); macrophages marked by F4/80 (red), M2 macrophages marked by CD163 (green) and dendritic cells marked by CD11c (blue).

**E)** UMAP embedding showing temporal dynamics for ECM scaffold clusters. Each dot is one ECM scaffold. Upon clodronate-treatment causing macrophage-depletion, macrophage-dominated ECM scaffold clusters appear delayed from normally regenerating tissues.

**F)** Heatmap showing the number of ECM scaffold cluster in uninjured and regenerating muscles at day 1, 3, 6, and 10 after injury with or without intramuscular injection with clodronate liposomes at day 2. n=4-8 per group.

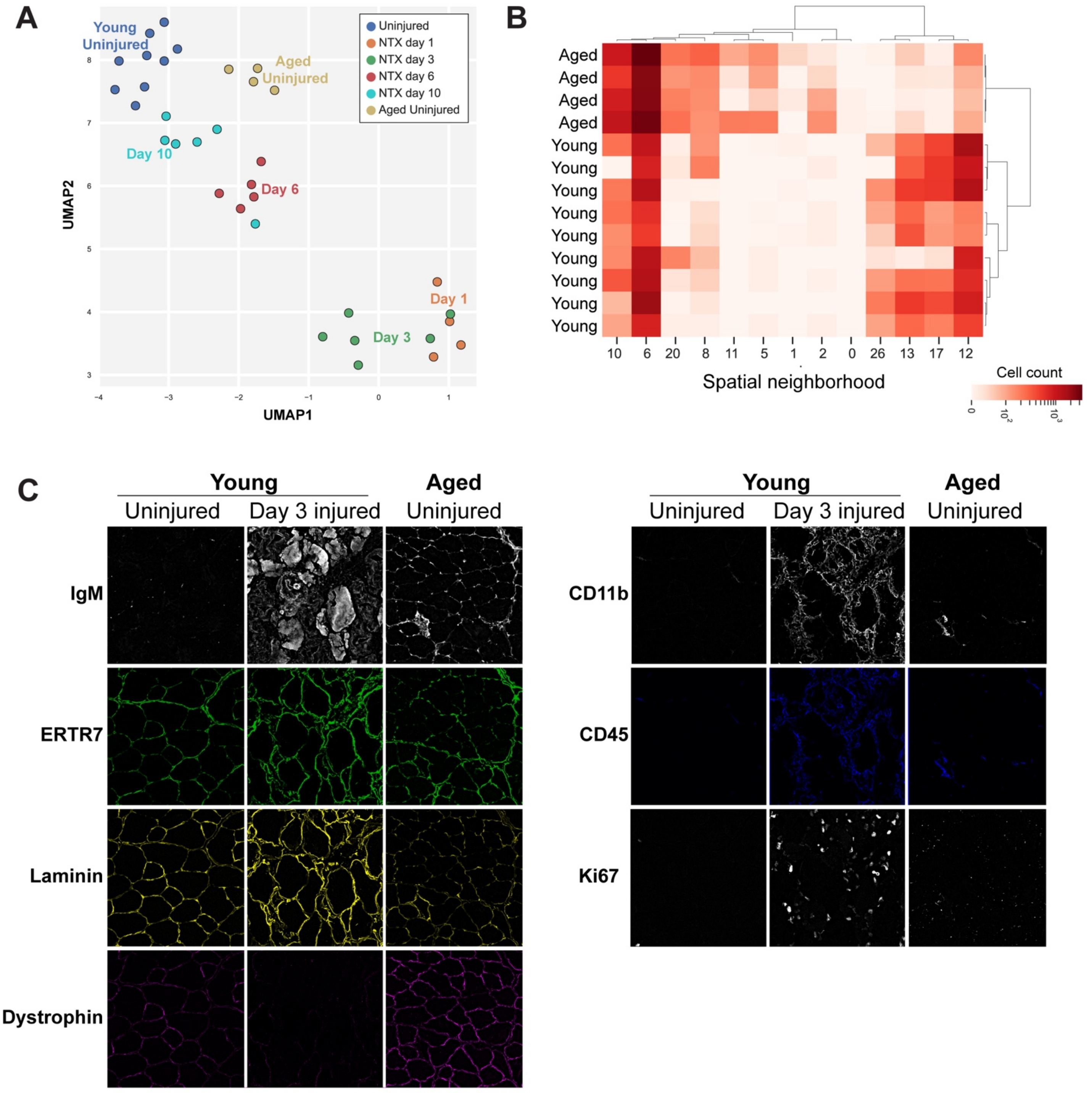


**Supplemental Figure S7. Cellular, molecular, and architectural alterations in aged skeletal muscle.** Related to Figures 5 and 6.

**A)** UMAP embedding of the cellular composition of uninjured and regenerating skeletal muscles from **Figure 2F** with aged uninjured muscle data. n=4-8 per group. Aged uninjured muscles cluster apart from young uninjured and day 10 regenerating muscles.

**B)** Heatmap showing cellular neighborhood dysregulation in aged muscle. Log transformed enrichment of spatial cellular neighborhood clusters from **Figure 4D** in uninjured muscles of young and aged mice. Each column is a biological replicate; n=8 young and 4 aged samples; Neighborhoods showing significant change ($p<0.05$) with aging are shown.

**C)** Aberrant localization of IgM in the ECM of aged muscle correlates to immune cell accumulation. Representative CODEX image of young uninjured muscle, day 3 after injury and aged uninjured muscle. IgM (gray, left), which accumulates in the injured myofibers after myotoxin injection, is found in the ECM (colocalizes with ERTR7 and Laminin; green and yellow, respectively) of aged uninjured muscles. Regions showing IgM accumulation also contain CD11b+ (gray, right top) and CD45+ (blue) immune cells. Ki67 (right bottom) marks dividing cells. The same field-of-view is shown in each column.
